# Molecular evidence of anteroposterior patterning in adult echinoderms

**DOI:** 10.1101/2023.02.05.527185

**Authors:** L. Formery, P. Peluso, I. Kohnle, J. Malnick, M. Pitel, K. R. Uhlinger, D. S. Rokhsar, D. R. Rank, C. J. Lowe

## Abstract

The origin of the pentaradial body plan of echinoderms from a bilateral ancestor is one of the most enduring zoological puzzles. Since echinoderms are defined by morphological novelty, even the most basic axial comparisons with their bilaterian relatives are problematic. Here, we used conserved antero-posterior (AP) axial molecular markers to determine whether the highly derived adult body plan of echinoderms masks underlying patterning similarities with other deuterostomes. To revisit this classical question, we used RNA tomography and *in situ* hybridizations in the sea star *Patiria miniata* to investigate the expression of a suite of conserved transcription factors with well-established roles in the establishment of AP polarity in bilaterians. We find that the relative spatial expression of these markers in *P. miniata* ambulacral ectoderm shows similarity with other deuterostomes, with the midline of each ray representing the most anterior territory and the most lateral parts exhibiting a more posterior identity. Interestingly, there is no ectodermal territory in the sea star that expresses the characteristic bilaterian trunk genetic patterning program. This suggests that from the perspective of ectoderm patterning, echinoderms are mostly head-like animals, and prompts a reinterpretation of the evolutionary trends that made echinoderms the most derived animal group.

## Introduction and results

Echinoderms, defined by their calcitic endoskeleton, unique water vasculature system, and perhaps most strikingly by their pentaradial body plan^1,2^, are among the most enigmatic animal phyla. Since echinoderms are phylogenetically nested within the deuterostomes^3-5^ (echinoderms, hemichordates, and chordates), their pentaradial organization was evidently derived from a bilaterian ancestor. Yet, despite a rich fossil record, comparative morphological studies have come to conflicting conclusions regarding the axial transformations that led to pentamery from the ancestral bilaterian state^2,6^. Among bilaterians, the deployment of the gene regulatory network that specifies ectoderm AP polarity is highly conserved and represents a suite of characters that is often more conserved than the body plans they regulate^7-14^ (Fig.1a). The deployment of this AP regulatory network could therefore provide an alternative way to test hypotheses of axial homology in cases such as echinoderms where morphological characters are too divergent to reconstruct ancestral states^15^.

**Figure 1:**
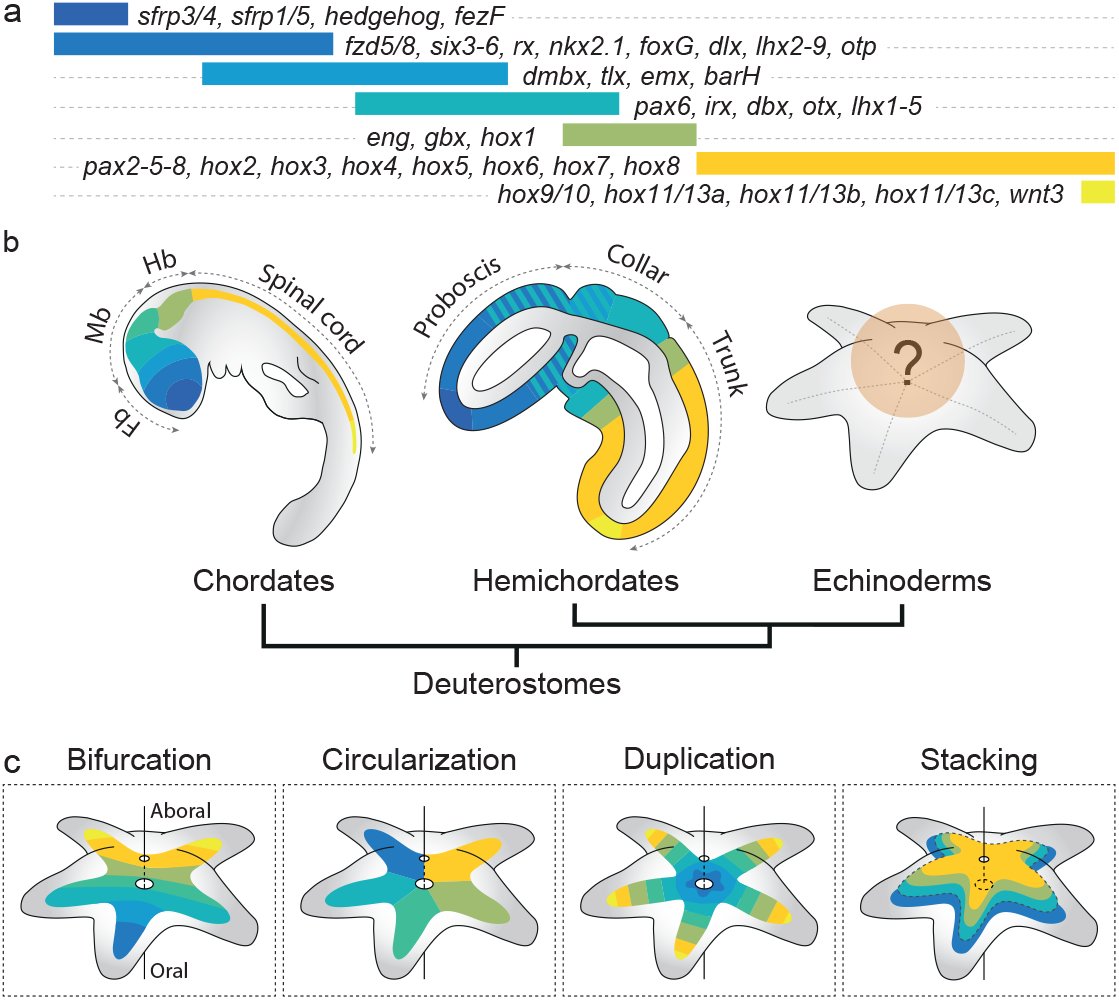
Deployment of the antero-posterior patterning system in deuterostomes. **a**, Expression map of the conserved transcription factors and signaling ligands involved in ectoderm patterning along the AP axis, as observed in the hemichordate *S. kowalveskii*. **b**, Previous work in chordates and hemichordates has demonstrated extensive regulatory conservation in ectodermal AP patterning, establishing the ancestral regulatory characteristics of early deuterostomes. How this system is deployed in echinoderms remains unclear. **c**, Four hypotheses have been proposed for the deployment of the AP patterning system in echinoderm adult body plan: bifurcation, circularization, duplication and stacking.

Detailed comparisons between chordate and hemichordate axial patterning have established the ancestral deuterostome AP patterning program^11-14^. The fate of this conserved network and its role in patterning the adult body plan of extant echinoderms could provide key insights into the evolution of echinoderm axial properties, with two distinct scenarios. First, the ancestral deuterostome AP patterning network could have been dismantled and reassembled into novel conformations during the radical body plan modifications along the echinoderm stem lineage. In this scenario, expression of transcription factors in derived morphological structures without conservation of relative spatial expression across the network would imply co-option into novel developmental roles^16,17^. Alternatively, conservation of spatially coordinated expression of this network during the elaboration of the echinoderm adult body plan would provide a molecular basis for testing hypotheses of axial homology with bilaterians, and establish regional homologies masked by divergent anatomies^15,18^ (Fig.1b).

Four main hypotheses have been proposed to relate the echinoderm body plan to other bilaterians (Fig.1c). The bifurcation^19^ and the circularization^19,20^ hypotheses can be ruled out, since they require a unique molecular identity for each of the five echinoderm rays that is inconsistent with molecular data^18,21^. In the duplication hypothesis^19,22^, each of the five echinoderm rays is a copy of the ancestral AP axis, and in the stacking hypothesis ^2,6,23,24^ the oral-aboral axis of adult echinoderms is homologous to the ancestral AP axis. While broad bilaterian comparisons of AP axis patterning are typically based on ectodermal expression domains, the stacking hypothesis was proposed largely on the basis of nested posterior Hox gene expression in the posterior mesoderm of the bilateral larval stages in holothuroids (sea cucumbers), crinoids (sea lilies), and echinoids (sea urchins)^25-31^, and need to be tested based on ectodermal expression.

### The transcriptional landscape of a pentaradial animal

We tested these classical hypotheses by examining gene expression using spatial transcriptomics and *in situ* hybridizations in the asteroid (sea star) *Patiria miniata* (Fig.2a,b). The manifestation of pentaradial symmetry in asteroids is simpler than in other echinoderm classes because it follows a planar organization with the rays extending around the oral-aboral axis through easily distinguishable alternating ambulacral and interambulacral territories (Fig.2b). This makes asteroids particularly well-suited to study the possible correspondence between bilaterian AP patterning genes and the pentaradial echinoderm body plan.

**Figure 2:**
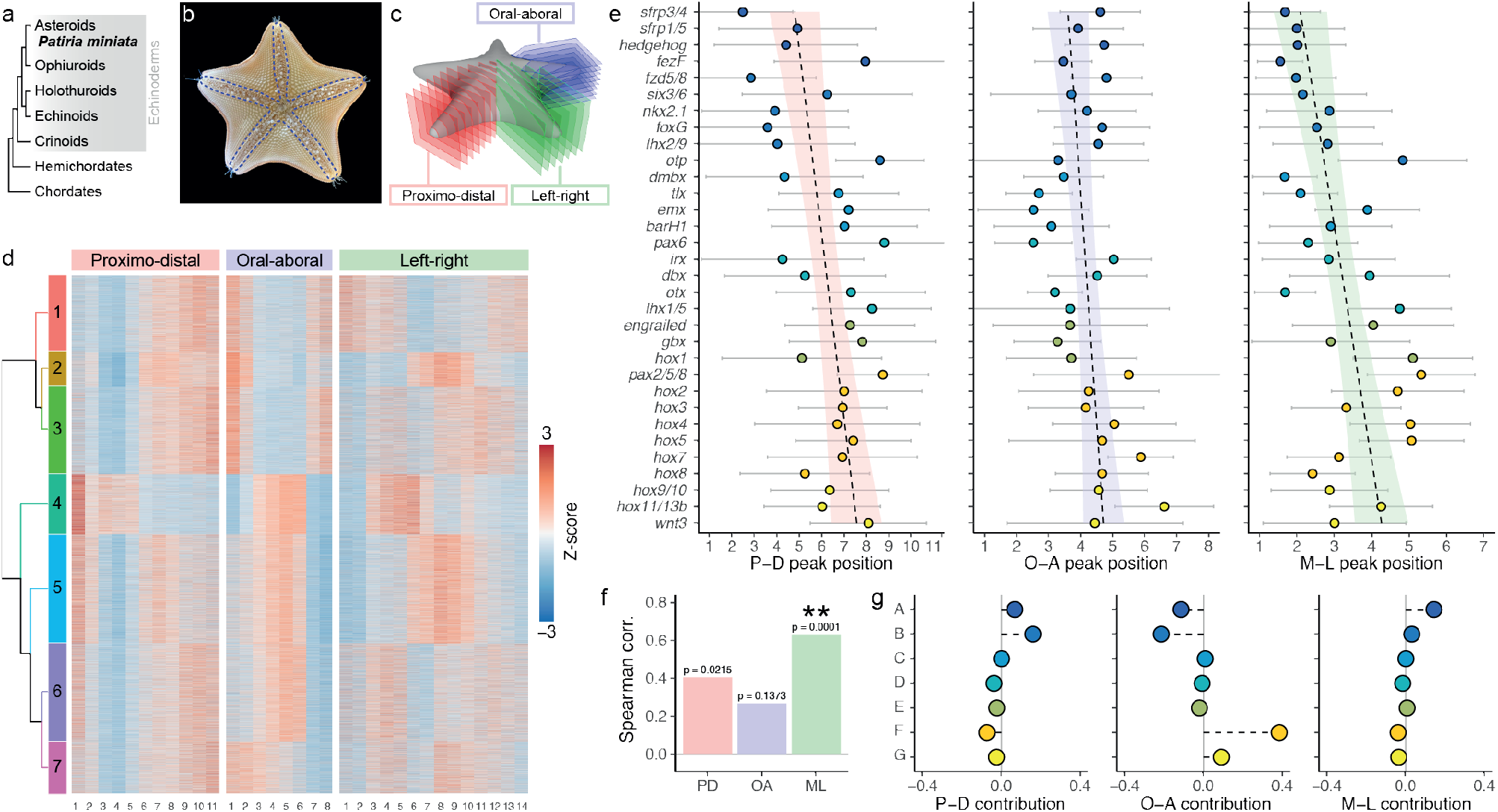
RNA tomography reveals medio-lateral dimension of the arms as the main driver of antero-posterior patterning system deployment in *Patiria miniata*. **a**, Phylogenetic position of *P. miniata* within deuterostomes, the grey box highlights the echinoderm phylum. **b**, Young *P. miniata* specimen, viewed from the oral side. The blue dotted lines outline the ambulacra. **c**, Experimental design of the RNA tomography. **d**, Gene expression z-scores along the P-D, O-A and L-R dimensions of the RNA tomography show 7 main trends of expression. **e**, Expression peak position of AP patterning related genes along the P-D, O-A and L-R dimensions. Genes are ranked from the most anterior to the most posterior based on their expression patterns in *S. kowalevskii*, with a linear model fitted to the peak positions (dotted lines). Error bars show the standard deviation of the probabilistic determination of the peak position. **f**, Spearman correlations between the antero-posterior ranking of the AP related genes and their peak position along the three dimensions of the RNA tomography. Only the M-L correlation is statistically supported (two-sided Spearman correlation test). **g**, Relative contribution of each group of AP patterning related gene to the Spearman correlation between the antero-posterior ranking of the AP related genes and their peak position along the three dimensions of the RNA tomography. Anterior gene groups drive the P-D and M-L correlations, while the Hox genes are the main drivers of the O-A correlation.

To investigate the transcriptional landscape of *P. miniata* along its body axes, we began with an unbiased spatial transcriptomics approach using RNA tomography^32^. We cryo-sectioned three arms from *P. miniata* young juveniles along three different dimensions: from the proximal to the distal part of the arm (P-D), from the oral to the aboral side (O-A), and from the left to the right side (L-R) (Fig.2c; Supplementary Fig.1). Sections were barcoded and pooled for single molecule real time sequencing with PacBio IsoSeq (Supplementary Fig.2a-e; Supplementary Tables 1-5), yielding a three-dimensional atlas of 25,794 gene expression profiles along the P-D, O-A and L-R dimensions of *P. miniata* arms (Fig.2d). Gene clustering based on the similarity of RNA tomography expression profiles highlighted seven principal patterns of gene expression (Fig.2d). We confirmed that the RNA tomography transcriptional landscape was consistent with the actual anatomy of the animal by considering Spearman correlations between sections along each dimension and analyzing the expression profile of marker genes known to be expressed in particular tissues (Supplementary Fig.3a-c). To aid in this analysis we generated a new *P. miniata* genome assembly (Supplementary Fig.2a,d,f; Supplementary Tables 5,6).

To consider possible molecular anatomical homologies across deuterostomes we identified 36 conserved molecular markers in the RNA tomography dataset that define specific ectoderm territories along the AP axis in hemichordates and chordates (Fig.1a; Supplementary Fig.4), and retrieved their expression profiles (Supplementary Fig.3d; Supplementary Table 4,7). These marker genes included transcription factors (*fezF, six3/6, nkx2*.*1, foxG, lhx2/9, otp, dmbx, tlx, emx, barH1, pax6, irx, dbx, otx, lhx1/5, engrailed, gbx, pax2/5/8, hox1, hox2, hox3, hox4, hox5, hox7, hox8, hox9/10, hox11/13b*); members of the Wnt signaling pathway (*sfrp1/5, sfrp3/4, fzd5/8, wnt3*); and the ligand *hedgehog*. Four additional transcription factor markers (*rx, dlx, hox11/13a* and *hox11/13c*) were excluded from the computational analyses because of low expression levels (Supplementary Fig.3e). *Hox6* is absent in our Hox cluster assembly, and was also missing from the crown of thorns sea star *Acanthaster planci*^33^, consistent with its loss prior to the divergence of these two asteroid families (Supplementary Fig.5).

The duplication and stacking hypotheses would be supported by staggered expression of these AP patterning markers along the P-D or O-A dimensions, respectively. We ranked the AP patterning markers from the anterior to posterior using as a template the hemichordate *Saccoglossus kowalevskii*, the most closely related bilaterian species with a comprehensive expression pattern dataset for these markers (Supplementary Fig.6a,b; Supplementary Table 7). We then tested the Spearman correlation between the ranking of the genes and the position of their peak of expression along the P-D and O-A dimensions (Fig.2e). In both cases, we found moderate but not statistically significant correlations (ρ=0.41, *p*=0.021 and ρ=0.26, *p*=0.13, respectively) (Fig.2f). After organizing the AP patterning genes into seven groups based on their expression profiles (Supplementary Fig.6b) we found that the moderate correlation with the O-A dimension was mostly explained by the Hox genes alone (Fig.2g). This was expected, since the stacking hypotheses was primarily informed by the sequential expression of Hox genes in mesoderm derivatives^6,23^.

Surprisingly, we found a much stronger correlation (ρ=0.63, *p*=1.4×10^−4^) between gene order and the medio-lateral axis (M-L) (Fig.2f). The most anterior genes appeared to be largely expressed close to the midline of the arm, while more posterior genes were expressed more laterally on either side of the midline. In this case, all the AP patterning marker groups had similar weight in the correlation (Fig.2g). To confirm that the observed correlations were not the result of ranking biases, we randomly shuffled the gene ranking within each group (Supplementary Fig.6c-e). For all the permutations, the M-L correlation was statistically supported, with a mean p-value of 3.8×10^−4^ and a median of 3.1×10^−4^, and was consistently superior to the P-D and O-A correlations (Supplementary Fig.6c-e). This suggests that neither the duplication nor the stacking hypotheses accurately describe the deployment of AP-related patterning genes in *P. miniata*, and that most of the underlying patterning logic of the pentaradial plan is not explained by existing models.

### Gene expression patterns suggest a medio-lateral deployment of the AP patterning system

RNA tomography gives average axial positional information across all germ layers. To further investigate axial patterning specifically in the ectoderm, we examined the expression pattern of the 36 marker genes using *in situ* hybridization chain reactions (HCR, Supplementary Fig.7; Supplementary Table 8) on post-metamorphic juveniles. This representative stage follows the resorption of larval structures into the rudiment during the course of the metamorphosis (Fig.3a). The elaboration of the pentaradial symmetry is initiated earlier within the larval mesoderm, but the mechanisms at play in this process are likely distinct from those that give their regional identity to the final axes of the pentaradial body. Accordingly, classical ectoderm AP patterning genes such as *pax6* and *otx* started to be expressed at the onset of metamorphosis but were most highly expressed in the post-metamorphic juveniles, when the definitive adult body plan is elaborated (Fig.3b,c). As in the adult, the juvenile body plan is organized into ambulacral territories on the oral side and interambulacral territories at the edge of the ambulacra and on the aboral side. The ambulacral ectoderm is divided in two main regions: the medial ambulacral ectoderm comprising the radial nerve cords (RNCs) and the circumoral nerve ring (CNR), and more laterally the epidermis covering the podia (Fig.3c,d). The remaining ectoderm is interambulacral and covers the lateral edges of the arms and the entire aboral surface of the adult.

**Figure 3:**
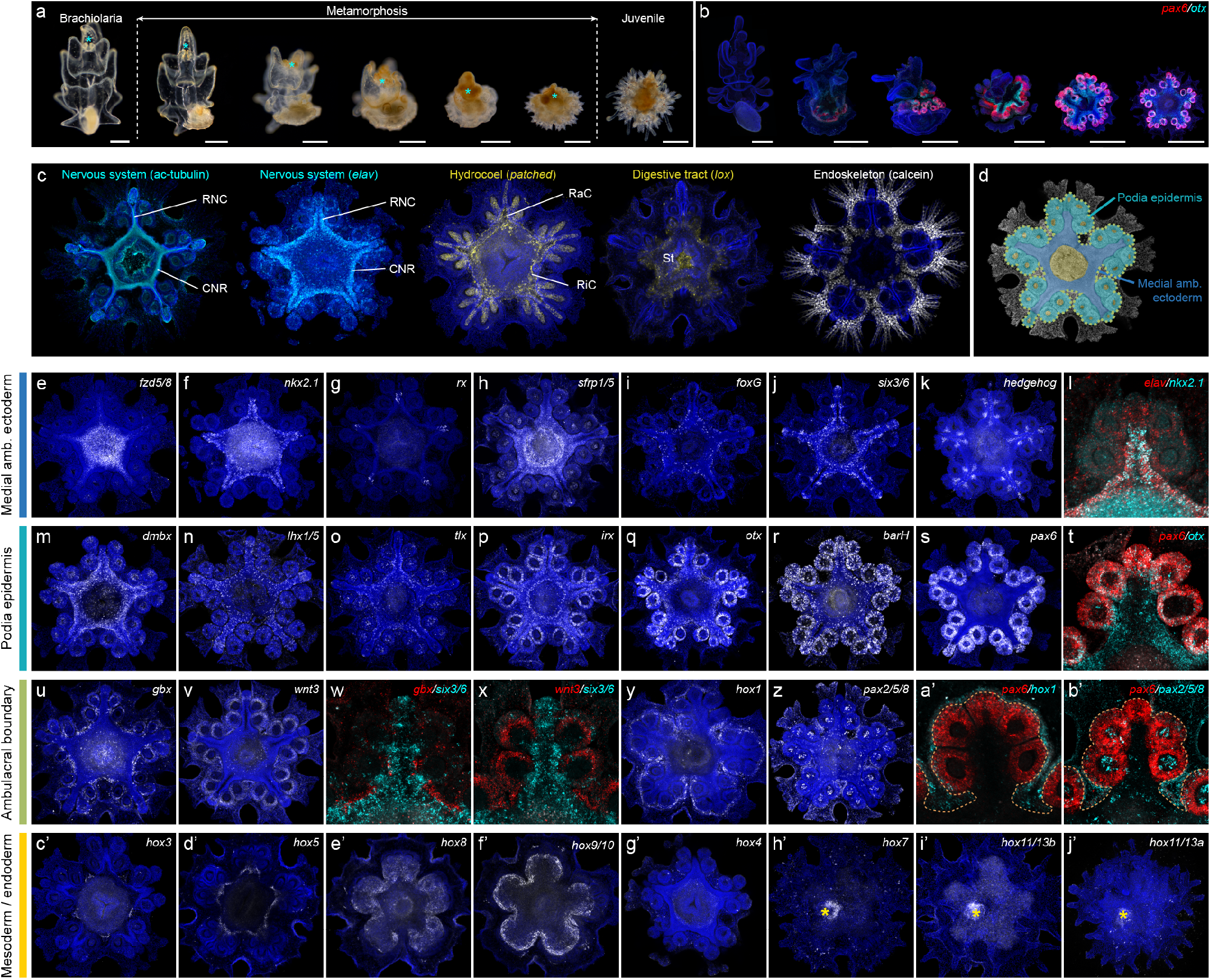
Gene expression data reveals the deployment of the antero-posterior patterning system in *Patiria miniata* ambulacral ectoderm. **a**, Metamorphosis in *P. miniata*. The anterior part of the brachiolaria larva (cyan asterisk) is resorbed into the rudiment. **b**, HCRs showing *otx* and *pax6* starting to be expressed during metamorphosis in the developing adult pentaradial body plan. In **a**,**b**, scale bar = 250μm. **c**, Stainings showing the main anatomical features of post-metamorphic juveniles, imaged from the oral side: the nervous system (cyan) stained with antibodies against acetylated-tubulin highlighting the nerve tracts and the neuronal marker *elav* highlighting the cell bodies, the hydrocoel stained with the marker *patched*, the digestive tract stained with the marker *lox* and the endoskeleton stained using calcein. RNC: radial nerve cords; CNR: circumoral nerve ring; RaC: radial canal; RiC: ring canal; St: stomach **d**, DAPI-stained (grey) specimen colored to highlight the main anatomical regions of the oral side of a post-metamorphic juvenile. The ambulacral ectoderm (outlined by the green dotted line) comprises two main regions: the medial ambulacral ectoderm (blue) and the podia epidermis (cyan). In some parts of the specimen internal germ layers are apparent through the confocal z-stack projection, such as the pharynx and the terminal ends of then hydrocoel (yellow). **e-j’**, HCRs of *P. miniata* juveniles imaged from the oral side (**e-g’**) or the aboral side (**h’**,**j’**). In **e-k**,**m-s**,**u**,**v**,**y**,**z**,**c’-j’** specimens are counterstained with DAPI (blue). **l**,**t**,**w**,**x**,**a’**,**b’** show a magnification on a single ambulacrum. **e-l**, Genes primarily expressed in the medial ambulacral ectoderm. Colocalization with elav indicates the expression in the CNR and the RNCs. **m-t**, Genes primarily expressed in the podia epidermis. **u-b’**, Genes primarily expressed in the ambulacral boundary. In **a’-b’** orange dotted lines outline the ambulacral ectoderm. **c’-f’**, Hox genes primarily expressed in the coeloms. **g’-j’**, Hox genes primarily expressed in the digestive tract. In **h’-j’** white asterisks indicate the position of the developing intestinal tract.

We considered the expression of four groups of patterning genes expressed along the AP axis in both hemichordates and chordates. The first group (*fzd5/8, nkx2*.*1, rx, sfrp1/5, foxG, six3/6, hedgehog*) has strong anterior ectodermal localizations in the proboscis of *S. kowaelvskii*^11,34,35^ and in the forebrain of vertebrates^13,14,36^ (Fig.1a,b). In *P. miniata* we found a similarly overlapping patterns of regional expression mostly restricted to the developing CNR, the RNCs, and in the case of *six3/6* and *hedgehog* at the edge of the RNCs at the level of each secondary podia (Fig.3e-l; Supplementary Fig.8a-e). This region corresponded to the most medial part of the ambulacral ectoderm (Fig.3d). These findings are inconsistent with the duplication hypothesis, which predicts a staggered expression of anterior to posterior markers along each RNC.

The next group (*lhx1/5, dmbx, tlx, irx, fezF, dbx, otx, barH* and *pax6)* overlaps with the previous anterior-class genes in *S. kowalesvskii* but with a more caudal distribution in the posterior proboscis and into the collar^11,34^; in chordates, these genes are primarily expressed either in the forebrain or the midbrain^13,14^ (Fig.1a,b). In *P. miniata*, expression of this group of genes overlaps in the most medial territory with the most anterior class genes, but with expanded lateral domains on either side of the RNCs (Fig.3d,m-s; Supplementary Fig.8f-h), into the epidermis covering the podia. *FezF* and *dbx* were only expressed in a limited number of cells in the ambulacral ectoderm (Supplementary Fig.9a,b). Only *pax6* was not expressed in the medial ambulacral ectoderm and was restricted to the podia epidermis, as previously reported in other echinoderm species^37,38^ (Fig.3t). Thus, the outer ambulacral ectoderm appears to have a more posterior molecular identity, similar to that of the hemichordate collar or the vertebrate midbrain.

The third category (*gbx, wnt3, hox1, pax2/5/8*) includes genes that have more posterior expression patterns in hemichordates and chordates (Fig.1a,b). *Gbx, hox1* and *pax2/5/8* are all expressed in hemichordates at the boundary between the collar and the trunk, and in chordates at the midbrain/hindbrain boundary, which delimits head and trunk identities^12,34,39,40^. *Wnt3* is expressed at the posterior end of the AP axis in both phyla^35,41^. In *P. miniata*, we found that these four genes are expressed at the boundary between ambulacral and interambulacral ectoderm (Fig.3d,u-b’; Supplementary Fig.8i-l). *Gbx* and *wnt3* were expressed in the outer part of the ambulacral ectoderm, establishing a mutually exclusive boundary with more anterior genes like *six3/6* (Fig.3u-x). *Hox1* and *pax2/5/8* were expressed more laterally compared to *gbx* and *wnt3*: *hox1* outlined the entire ambulacral area, while *pax2/5/8* had a more complex expression pattern and was primarily expressed between the ambulacra (Fig.3y-b’). We suggest that in *P. miniata* these genes marked the outer limit of an anterior compartment.

Finally, Hox genes are expressed in hemichordates and chordates posteriorly to the collar/trunk and midbrain/hindbrain boundary, respectively, and are involved in trunk patterning^11,39,42^ (Fig.1a,b). In *P. miniata*, only *hox1* was detected in the ectoderm. *Hox3, hox5, hox8* and *hox9/10* were expressed in mesoderm derivatives (Fig.3c’-f’). *Hox4* was barely above the detection threshold but was expressed in the pharynx (Fig.3g’), while *hox7, hox11/13a* and *hox11/13b* were expressed in the developing intestinal tract (Fig.3h’,j’). These observations agree with previous reports that Hox expression in echinoderms is largely restricted to internal germ layers^25,26,29,30^ and with the hox-driven O-A correlation observed in our RNA tomography dataset. In hemichordates and other bilaterians, Hox gene domains are intercalated during trunk development between the anterior domains and the posterior end of the animal, which expresses posteriorizing factors such as *wnt3*^39,43^. In *P. miniata*, we suggest that there is no ectoderm equivalent to a trunk region, because *wnt3* is expressed at the edge of the ambulacral region, and because *hox1* is the only Hox gene expressed in the ectoderm. Therefore, the deployment of the AP patterning system in *P. miniata* seems to be limited to the ambulacral region and its boundary.

Not all genes shared the same conserved relative expression than in hemichordates and vertebrates, either because they were not detected (*dlx*), not expressed in the ectoderm (*engrailed, sfrp3/4*), or exhibited different relative spatial arrangements (*lhx2/9, emx, otp*) (Supplementary Fig.9c-h). This presumably reflects the plasticity of the AP patterning system and its adaptation to the radically different pentaradial body plan. Despite these discrepancies, germ-layer specific expression patterns corroborate that the M-L dimension of the RNA tomography, and not the O/A or P/D, is the main and unexpected driver of the ectoderm AP patterning logic of the pentaradial body plan in *P. miniata*.

### Evolution of axial properties in echinoderms

The organizational modifications to the ancestral bilaterian deuterostome body plan during early echinoderm evolution were so profound that even basic axial comparisons with other deuterostome taxa are problematic at the morphological level^44,45^. Here we investigated this extensive body plan reorganization by spatially mapping the deployment of the ancestral bilaterian ectodermal AP patterning system in the pentaradial body plan of *P. miniata*. Since this patterning system is largely conserved between hemichordates and chordates, we can confidently reconstruct its ancestry in early deuterostomes and at the base of the ambulacrarians^11-14^.

We find that that much of the ancestral anterior patterning network is spatially deployed in a manner incompatible with previously proposed hypotheses of echinoderm axial homologies^6,18,19^. Rather, expression patterns map onto a novel coordinate system that we call the ambulacral-anterior model of echinoderm body plan evolution (Fig.4). In this model, the midline of each ambulacrum expresses the most anterior bilaterian molecular identity, equivalent to the forebrain and proboscis in vertebrates and hemichordates, respectively. The mid-lateral regions on either side of the nerve cords, including the ectoderm wrapping the podia, share patterning similarities with more caudal ectodermal territories of hemichordates and chordates, down to the collar and midbrain, respectively. Finally, the ambulacral boundary at the edge of the ambulacral ectoderm displays the most posterior molecular profile corresponding to the collar/trunk boundary of hemichordates and the midbrain/hindbrain boundary of vertebrates. The ambulacral-anterior model predicts that echinoderms are the first example of bilaterians in which the “anterior” identity is located at the center of a sheet of tissue, rather than being located at an extremity. In addition, despite their organizational disparity, the anatomical output of the anterior-like domain, such as neural condensations and an extensive array of sensory structures, shares similarities across all deuterostome phyla, including echinoderms.

**Figure 4:**
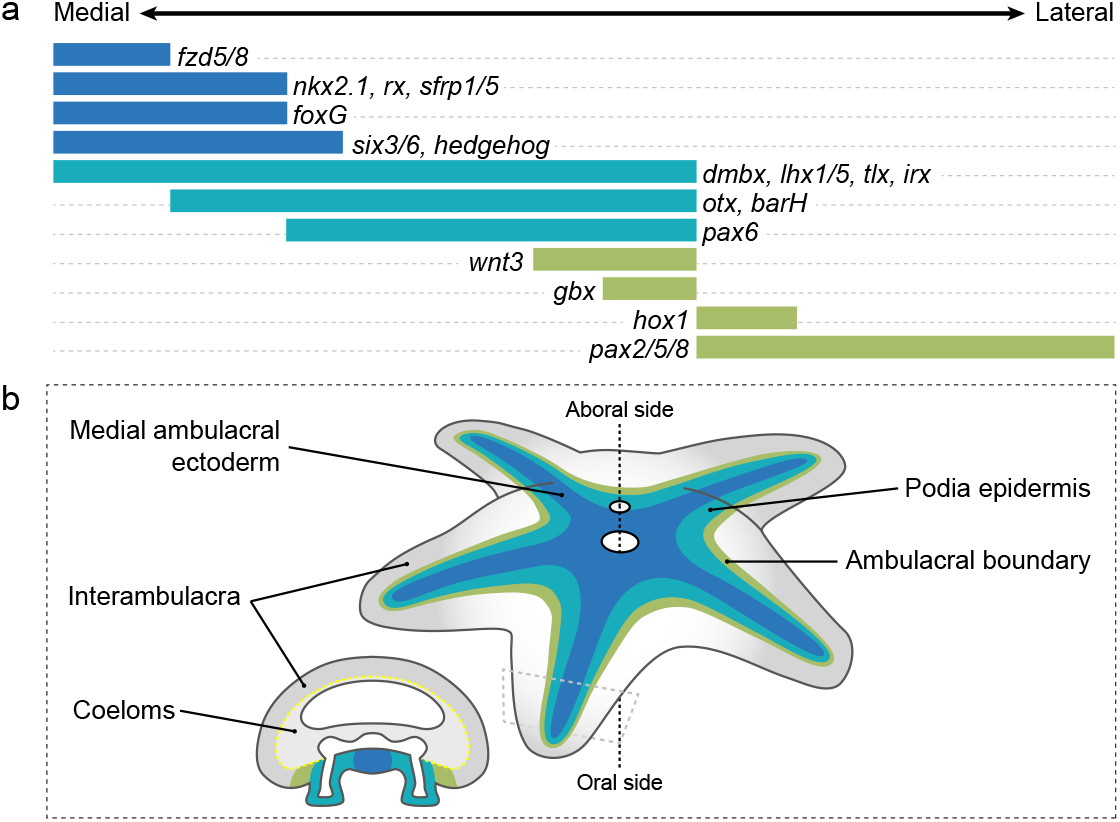
Ambulacral-anterior model of echinoderm body plan evolution. **a**, Expression map of the conserved transcription factors and signaling ligands involved in the ambulacral ectoderm patterning in *P. miniata* and organized from the midline of the ambulacrum (left) towards the interambulacrum (right). **b**, Diagram of the ambulacral-anterior model in a generalized asteroid with a cross-section through one of the arm. Only genes expressed in the ectoderm are shown.

Strikingly, despite the presence of a genomic Hox cluster, ectodermal Hox gene expression is largely absent (except for *hox1*), suggesting a loss of the ancestral ectodermal trunk regulatory program. Yet, Hox genes are expressed in the mesoderm and endoderm, displaying a marked uncoupling of germ layer AP patterning as proposed by Lacalli^46^. The mesoderm and endoderm trunk programs are surrounded by an ectoderm with two clear territories, including (1) an ambulacral anterior-like domain that is the main focus of our work, and also surprisingly (2) an extensive interambulacral domain that wraps around the aboral side of the animal and displays uncertain axial identity, without coherent deployment of the ancestral AP patterning program. The uncoupling of an ectodermal head and trunk programs is not unique to *P. miniata* and has been demonstrated in both larval echinoderms and hemichordates^39,47^, and recently in annelid larvae^43^, suggesting that these regulatory programs can be uncoupled over macroevolutionary time frames.

It could be argued that the bilaterian-like relative spatial expression of AP markers found in *P. miniata* is the result of co-option rather than modification of an ancestral AP patterning system. However, co-option of gene regulatory networks into the development of novel structures usually involves a limited number of genes^48,49^; there are to our knowledge no examples of co-option of the entire anterior part of the AP patterning program. Our finding that 18 out of the 22 patterning genes carrying an anterior identity are consistently deployed in the ambulacral ectoderm during the development of *P. miniata* juveniles, however, suggests that previous proposals that co-option of individual transcription factors is linked to the emergence of the derived pentaradial body plan^16,17^ are unlikely.

Did the reorganization of the ancestral AP axis documented here occur during the early stem evolution of echinoderms, or did it occur during the latter diversification of asteroids? So far, our model is consistent with published findings from two other major echinoderm clades, crinoids and echinoids. In crinoids, the expression of *six3/6, otx*, and *pax6* has been reported in the ambulacral ectoderm^50^. In echinoids, AP patterning related genes have been extensively surveyed in *Peronella japonica*^24,30^ and to a lesser extent in other species^25,37,38^. In both cases, most of the anterior patterning related genes studied show expression patterns compatible with our model. Furthermore, Hox gene expression has been described in echinoids, holuthuroids, and crinoids^25-31^ and is also largely congruent with the expression patterns that we observed in *P. miniata*. This suggests that our observations reflect early regulatory changes that occurred during stem echinoderm evolution. Further comprehensive comparative analyses will be required to test this hypothesis.

The evolution of body plan variations across echinoderm classes has proven challenging to reconstruct based on morphological features alone^6,44^. Our model offers a powerful tool to establish robust regional homologies between classes and an independent dataset to test existing competing hypotheses. Most importantly, the new axial paradigm established here can be integrated with the exquisite fossil record of the phylum to reinvestigate key morphological transformations in light of regulatory changes.

## Acknowledgements

The authors would like to thank Fabio Benedetti for the help with the RNA tomography analyses, Andrea Dingeldein for the anatomical illustrations, Antoine Formery for the 3D models of the RNA tomography sections, Auston Rutledge for helping with animal husbandry, and Veronica Hinman for providing clones for colorimetric *in situ* hybridizations. The authors also thank Gregory A. Wray, Thurston Lacalli, Jeffrey R. Thompson, Rich Mooi, Jenifer C. Croce, and the members of the Rokhsar and Lowe laboratories for discussions. This work was funded by a NASA grant to C.J.L (NNX13AI68G), a NSF grant to C.J.L (1656628) and Chan Zuckerberg BioHub funding to D.S.R. and C.J.L.

## Data availability

All data are available upon request.

## Code availability

Custom code used for RNA tomography analyses is available upon request.

## Authors contribution

Experiments were designed by L.F, P.P, D.R.R and C.J.L. Preliminary data were acquired by I.K., J.M. and K.R.U. Genome sequencing was done by P.P. and D.R.R. RNA tomography sectioning and sequencing was done by P.P, M.P, D.R.R and C.J.L. RNA tomography analyses were performed by L.F and D.S.R. Immunofluorescence, HCRs and imaging were performed by L.F. Data were analyzed by L.F, D.S.R and C.J.L. The manuscript was written by L.F, D.S.R, and C.J.L with inputs from all authors.

## SUPPLEMENTARY INFORMATIONS

## Supplementary Methods

### Animal husbandry and tissue preparation

Adult and young (∼2cm wide) *Patiria miniata* specimens were collected off the coast of Monterey bay, California, US, and kept in circulating sea water tanks. On one hand, to generate RNA tomography datasets and genomic DNA (see below), two young specimens (referred to as specimens #1 and #2) were used. They were anesthetized in filtered sea water and 7.5% MgCl (1:1), and then four arms were dissected out (Supplementary Fig.1a,b). One of the arm was proceeded for DNA extraction, and the three others were included in HistoPrep embedding medium (ThermoFischer). On the other hand, to generate fixed material for HCRs and immunochemistry (see below), several batches of gravid adults were spawned by injecting 1ml of 1μM 1-methyladenine (Acros Organics) in each of the gonads. Sperm and mature oocytes were released by the animals about 45’ and 90’ after the injection, respectively. Following *in vitro* fertilization, embryos were cultured at 14°C in UV-sterilized filtered seawater, first at a density of about 100 embryos per mL and within 3L glass jars oxygenated by a motorized paddle. At 48 hours post fertilization, the culture concentration was adjusted to about 1 larva per mL, and from that point about 90% of the seawater was renewed every 2 or 3 days. Following water renewal, the larvae were fed *ad libidum* with freshly grown *Rhodomonas lens* microalgae. Brachiolaria larvae started to settle on the glass jars and to undergo metamorphosis between 1 and 2 months post fertilization. After metamorphosis, juveniles were collected from the glass jars by being first relaxed in a 1:1 mix of 7.5% MgCl2 and seawater and then being gently detached using a paintbrush.

### Genomic DNA isolation

Genomic DNA (gDNA) for genome sequencing was isolated from a dissected arm of the young *P. miniata* specimen #1. Using an extended handle conical tip pestle (Bel-Art Proculture) the *P. miniata* arm was homogenized in the presence of the extraction buffer and of proteinase K. Genomic DNA was then isolated using the DNeasy Blood and Tissue Kit (Qiagen) following the manufacturer’s instructions.

### Genome sequencing

#### Ultra-Low Input HiFi library preparation

Using gDNA from the arm of specimen #1, we generated a HiFi genome of *P. miniata*. The general workflow is described in Supplementary Fig.2a. As the gDNA isolated from the arm was predominantly shorter than the 10-15 kb which is recommended size for HiFi genomic library creation, a size selection was performed prior to doing an Ultra-Low Input (ULI) amplification and library preparation to remove fragments <7kb. The size selection was done on a SAGE BluePippin system using the 0.75% Agarose Dye-free Gel Cassette and the S1 Marker (SAGE). Approximately 100ng of DNA was recovered post size selection and used as input for the ULI PCR-based HiFi library protocol. The sample was amplified using the SMRTbell gDNA Amplification Kit (PacBio) and HiFi SMRTbell library was constructed using the SMRTbell Express Template Prep Kit 2.0 (PacBio) following manufacturer’s recommended protocol. After library construction, a final size selection was performed on the SAGE BluePippin as previously described using a size cut-off of 7kb. Library size was characterized on an Agilent 2100 BioAnalyzer using the DNA 12000 kit (Agilent). The additional size selection ensured having a final library with a fragment size range greater than 7kb.

#### HiFi reads sequencing

Sequencing reactions were performed on the PacBio Sequel II System with the Sequel Sequencing Kit 2.0 (PacBio). The kit uses a circular consensus sequencing (CCS) mode which provides >99.5% single molecule read accuracy^51^. The samples were pre-extended without exposure to illumination for 2 hours to enable the polymerase enzymes to transition into the highly progressive strand-displacing state and sequencing data was collected for 30 hours to ensure maximal yield of high-quality CCS reads. CCS reads were generated from the data using the SMRT Link Version 9.0 (PacBio). For the genomic HiFi sequencing, the library was bound to the sequencing enzyme using the Sequel II Binding Kit 2.2 and the Internal Control Kit 1.0 (PacBio). The HiFi reads generated 15,614,751 HiFi reads with a mean read length of 9,039 bp ± 1,671bp (Supplementary Fig.2f).

#### Genome assembly

Prior to *de novo* genome assembly, the reads were trimmed on the ends to remove any PCR primer sequences from the ultra-low amplification process using lima v.2.2 (PacBio). The forward sequence of the amplification adapter used was AAGCAGTGGTATCAACGCAGAGTACT. Once the HiFi reads were trimmed, a filtering step was performed to remove duplicate reads from the PCR step using SMRT link v.9.0. After removing duplicate reads, the HiFi reads (∼100X genomic coverage) were used to generate a draft diploid assembly using Hifiasm v.0.15^52^. This resulted in two highly contiguous haplotype primary assemblies of 680 Mb and 674 Mb respectively (Supplementary Table 6). Assembly completeness was assessed with the Benchmarking Single Copy Ortholog^53^ (BUSCO v.3.0) gene set for Metazoa at 94.8% for each of the individual haplotype primary assemblies and 96.2% overall for the diploid assembly as a whole (Supplementary Fig.2d; Supplementary Table 5). The two haplotypes have been deposited at DDBJ/ENA/GenBank under the accession numbers JAPJSQ000000000 and JAPJSR000000000.

### RNA tomography section preparation and RNA extraction

The three arms embedded in HistoPrep medium were cryosectioned using a Leica CM3050-S microtome along the appropriate dimension: proximo-distal (P-D), oral-aboral (O-A) or left-right (L-R) (Supplementary Fig.1b,c). The blocks used for the P-D and O-A dimensions came from the specimen #1, while the block used for the L-R dimension came from the specimen #2. Slice thickness was set to 25μm for the P-D and O-A dimensions and to 30μm for the L-R dimension, resulting in total in 430, 160 and 160 slices for the three respective dimensions. While sectioning the blocks, every 20 (P-D, O-A) or 10 (L-R) contiguous slices were pooled together into 1.5mL tubes. These resulted in a total of 22 tubes for the P-D dimension, 8 tubes for the O-A dimension and 16 tubes for the L-R dimension. Each tube was then processed for RNA extraction using a modified Trizol/RNeasy RNA extraction protocol^54^. In each tube the slices were homogenized in 1mL of Trizol using an extended handle conical tip pestle (BelArt Proculture). After vigorously mixing the Trizol homogenate with chloroform, each tube was centrifuged at 10,000xg RCF for 18’ at 4° C. The aqueous phase containing the RNA was carefully removed and the RNA was further purified using the RNeasy Plus Micro Kit (Qiagen) following the manufacturer’s instructions.

### RNA tomography

#### Barcoded cDNA IsoSeq SMRTbell library preparation

Using RNA isolated from the three sets of cryosections, we generated a RNA tomography^32,55^ dataset for *P. miniata* juveniles. The general workflow is described in Supplementary Fig.2a. Barcoded PacBio IsoSeq SMRTbell libraries were constructed using the SMRTbell Express Template Prep Kit 2.0 (PacBio) following the manufacturer’s instructions. We used a set of 22 barcoded sequences (Supplementary Table 1) which were combined with each RNA extracts (Supplementary Table 2). Typically, 12-14 PCR amplification cycles were used to generate enough barcoded double-stranded cDNA for the library preparation and subsequent sequencing runs (Supplementary Table 2).

#### Library sequencing

For the IsoSeq transcript libraries, library was bound to the sequencing enzyme using the Sequel II Binding Kit 2.1 and Internal Control Kit 1.0 (PacBio) and the sequencing reactions were performed on the PacBio Sequel II System. The three dimensions were sequenced independently, since there were shared barcodes between the different libraries. We obtained a total of 71,582,642 reads (Supplementary Table 3) with a mean read length of 3,843 bp, 3,152 bp and 2,450 bp for the P-D, O-A and L-R dimensions, respectively (Supplementary Fig.2b). In addition, HiFi read length distributions were consistent across each barcode within the respective RNA tomography dimension (Supplementary Fig.2c). Sequence read archives were deposited at DDBJ/ENA/GenBank under the bioproject PRJNA873766 and individual accession numbers for each barcode are provided in Supplementary Table 3. Recent advancements in full length transcript concatenation protocols^56^, as well as higher multiplexed SMRT sequencing flow cells have increased throughput with a concomitant decrease in sequencing cost over a single 24 hour run. Genomic, epigenetic and transcript data sets can now all originate from the same individual greatly improving the mappability and subsequent analysis of transcripts in highly polymorphic non-model organisms where gene sequences can differ by >5% in coding regions and as much as 40% in UTR sequence which complicate short read transcript alignment and reliable quantitation.

#### IsoSeq reads demultiplexing and refining

For each HiFi read file generated, the data was demultiplexed into barcode specific read files using lima v.2.2 (PacBio) and the barcodes listed in Supplementary Table 1. Once the data was demultiplexed each read file was refined to include only full length non-chimeric reads using isoseq3 v.3.4.0^57^. Chimeras were identified by inclusion of 5’ or 3’ RT-PCR primer sequences internal to the initial ‘full length’ HiFi read. The primer sequences used were NEB_5P (GCAATGAAGTCGCAGGGTTGGG), Clontech_5P (AAGCAGTGGTATCAACGCAGAGTACATGGGG), and NEB_Clontech_3P (GTACTCTGCGTTGATACCACTGCTT). Transcript clusters were identified using Cupcake v.25.2.0^58^ which leverages a genomic reference alignment-based strategy to identify redundant isoforms/transcripts for gene loci. This process was performed using each haplotype of the full diploid genome assembly of the same individual that the cDNA sequences were derived. The process of using the identical individual specimen for both genomic and transcriptomic datasets simplified the identification of transcripts in a highly polymorphic organisms such as *P. miniata*, by ensuring highest concordance of transcripts. We leveraged the diploid assembly to minimize dropouts that might be caused by haplotype specific null alleles and/or poor mapping between haplotypes for the same locus. Comparative alignment of the two haplotype derived transcriptomes facilitated further collapse of the gene set to one representative per loci.

#### Transcriptome generation and curation

Using each haplotype primary assembly, the complete set of full length non-chimeric transcript reads (FLNC) were clustered and collapsed to reduce gene redundancy while maintaining the highest possible level of gene completeness. The complete set of FLNC reads across all three combined RNA tomography datasets were clustered from a list of input bam files based on each of the haplotype specific primary assemblies. Minimap2 v.2.21^59^ was used to align the FLNC reads to each primary assembly and Sqanti2 v.7.4^60^ was used to cluster and filter redundancies. Once minimal transcript sets were obtained for each haplotype, a comparative alignment between the sets was performed using Minimap2 v.2.21 aligner to find unique transcripts between the two. From the ouput paf files, transcripts unique to haplotype 2 which were not present in the haplotpye 1 were identified, filtered out from the haplotype 2 set, and added to the Haplotype 1 set to obtain a more complete single copy transcript set.

Manually curated developmental genes of interest (Supplementary Table 7) were identified (see below, orthologues identification). All sequence in our “single copy” transcript set were aligned using TBLASTX to the manually curated set to (1) remove any duplication of these key transcripts in our reference transcriptome (e.g. the high polymorphism rate resulted in both copies of some of the manually curated genes being present) and (2) remove partial duplications (non-full length transcripts matching the manually curated set). The result of this curation step was to ensure that for every gene of interest, we had only one sequence in our reference transcriptome to maintain accurate downstream quantitation. Given the occasional duplication of transcripts in our reference transcriptome (eliminated for our genes of interest), it is assumed that the un-curated transcripts have some level of duplication.

Once we derived a near complete curated transcriptome for *P. miniata*, the barcode specific demultiplexed RNA tomography datasets were aligned to the transcriptome reference using Minimap2 v.2.21. The final refined *P. miniata* transcriptome comprised 25,794 transcripts and represented a nearly complete (91.5%) set of metazoan BUSCO genes (Supplementary Fig.2d, Supplementary Table 5). Transcript expression counts for each of the 25,794 gene model in our reference transcriptome were tallied using a simple Perl v.5.30.1 script that only counted primary alignments (no supplementary alignments) with a quality value of 15 or greater. Each section was tallied independently and the data merged. The alignment counts for each section were normalized to the total reads for each barcode in order to allow spatial comparisons to be made across each dimension of the RNA tomography dataset. This was done to account for variable recovery of total RNA in each barcoded section. (Supplementary Fig.2e).

### Orthologues identification

*P. miniata* orthologues of developmental genes of interest (Supplementary Table 7), which included 36 AP patterning related genes, the pan-neuronal marker *elav*, the gut marker *lox* and the hydrocoel marker *patched* were identified from the FLNC reads by reciprocal best blast hit and validated by phylogenetic trees (Supplementary Fig.4). Nucleotide sequences for these transcripts were deposited at DDBJ/ENA/GenBank and accession numbers are provided in Supplementary Table 7. Trees were calculated with both the Maximum likelihood and Bayesian inference methods. Maximum likelihood trees were calculated in MEGA v.7.0.26^61^ with the robustness of each node being estimated by bootstrap analyses (in 1000 pseudoreplicates). Bayesian inference trees were calculated using MrBayes v.3.1.2^62^ in 1,000,000 generations with sampling of trees every 100 generations and a burn-in period of 25%. The branching pattern of the ML tree was retained in the final tree figure, displaying, at each node, the bootstrap support of the ML analysis as well as the posterior probability support of the BI analysis.

### RNA tomography analyses

RNA tomography analyses were performed with R v.4.1.2 using custom-written code. For downstream analyses, the 22 sections of the P-D were merged pairwise by simple addition of the read counts mapping for each transcripts, bringing the final number of sections in the P-D dimensions to 11. In addition, section 14 of the L-R dataset was removed because it yielded a total read count 77% lower than the average total read count per section for the L-R dimension, which could have biased the quantification analyses. To maintain the symmetry in the L-R dimension, section 3 was also removed, bringing the final number of sections in the L-R dimension to 14.

Individual read counts from each section of the three RNA tomography dimensions were normalized against the total read count of the section in order to account for sequencing depth differences between the three dimensions and for geometrical disparities between the different sections of a single dimension. Because we were interested in the profile of highly expressed and variable genes, whereas genes with a uniform expression across the sections were poorly informative, a cutoff was applied to discard genes which consistently had the 20% lowest average expression level or 20% lowest variability in all of the three dimensions, resulting in a final set of 21,847 gene models (Supplementary Fig.3e). Of the 36 AP patterning related genes investigated in this study, four of them fell below the cutoff and were excluded from the computational analyses because of their low expression levels: *rx, dlx, hox11/13a*, and *hox11/13c* (Supplementary Fig.3e). For further analyses, the expression levels of each gene along each of the three dimension sections was transformed into z-score. For the clustering analysis, the dimensionality of the dataset was first reduced using a principal component analysis (PCA) performed simultaneously on the three dimensions of the RNA tomography. Kaiser-Guttman’s criterion^63^ was used to select the significant PCs of the PCA. Then, the coordinates of the transcripts along the retained PCA axes were used to compute the Euclidean distance matrix and hierarchical agglomerative clustering using Ward’s aggregation to produce an expression profile dendrogram^64^. The number of clusters was determined using a silhouette index^65^ and on biological significance.

We determined the ranking of the 36 investigated AP patterning related genes based on their expression profiles in the hemichordate *S. kowaleskii*, which is the closest bilateral echinoderm-relative in which the entire AP patterning program has been previously described^11,12,34,35, 66-69^. Gene expression pattern in other hemichordate studies are largely consistent with those observed in *S. kowalevskii*^39^. The 36 genes were divided into seven groups based on their expression domains (anterior proboscis (A), proboscis (B), anterior collar (C), posterior proboscis and collar (D), collar-trunk boundary (E), trunk (F), posterior tip of the trunk (G)) (Supplementary Fig.6a,b). Within each group, genes where ranked based on their expression profile and when required tied based on the available gene expression patterns in other closely related species.

To test the correlation between the AP patterning related gene ranking and the gene expression profiles along the three dimensions of the RNA tomography, we summarized the expression profiles to the position of a single expression peak to account for ambiguities with some of the genes that had a multimodal expression profiles. The peak position for each gene and in each dimension was determined probabilistically, using the expression z-score as a probability law over 100 replicates. The results of the replications was then averaged to give the peak position. The peak position was then correlated with the gene ranking using Spearman rank correlation. The statistical significance of the correlation was assessed by a two-sided Spearman test with a *p*-value threshold of 0.01. Furthermore, we assessed the contribution of each of the seven gene groups by calculating the difference between the correlation with and without each one of the seven groups. We were confident that the assignment of each gene to the groups A to G based on their expression profile in *S. kowalevskii* was robust to interpretation biases (Supplementary Fig.6a,b). On the other hand, we recognize that the internal gene ranking within each group was more subject to possible interpretation biases. To ensure that the observed correlations were not the result of such biases, we simultaneously shuffled the ranking of the gene within each group 100,000 times, and calculated the range of correlations and associated *p* values. The M-L correlation appeared consistently higher and more statistically supported than the P-D and O-A correlations, indicating that this was robust to any ranking bias (Supplementary Fig.6c,d). In addition, we determined the variability of the range of correlation values that was due to each group of gene by shuffling the ranking within each of group of gene independently 1,000 time. This showed that group D (genes expressed in the posterior proboscis and collar), which we assessed as the most ranking biases-prone, actually had little impact on the correlation range, compared to group B (genes expressed in the proboscis) and group F (genes expressed in the trunk, in particular Hox genes) which were less susceptible to ranking biases (Supplementary Fig.6e).

### *In situ* hybridization

Short *in situ* hybridization antisens DNA probes were designed based on the split-probe design of HCR v.3.0^70^ using HCR 3.0 Probe Maker^71^ with adjacent B1, B2 or B3 amplification sequences depending on the genes (Supplementary Table 8). At least 14 probe pairs were designed for each gene including the CDS, and for some of them adjacent 5’ and 3’ UTRs. The probe pairs were then ordered as oligo pools (Integrated DNA Technology) and suspended in nuclease-free water at a concentration of 0.5μM.

*In situ* hybridization were performed as outlined in Choi et al.^70^. *P. miniata* brachiolaria larvae, metamorphosing larvae and juveniles were incubated in fixation buffer (1X phosphate buffered saline (PBS), 0.1M MOPS, 0.5M NaCl, 2mM EGTA, 1mM MgCl2) containing 3.7% formaldehyde overnight at 4°C. Fixed samples were then dehydrated in methanol and stored at −20°C for at least 24 hours. The samples were progressively rehydrated in PBS containing 0.1% Tween-20 (PBST). They were permeabilized in detergent solution (1.0% SDS, 0.5% Tween-20, 150mM NaCl, 1mM EDTA (pH 8), 50mM Tris-HCl at pH 7.5) 30’ for larvae and 2 hours for juveniles. For juveniles, this was step was followed by an extra permeabilization in 4μg/mL proteinase K (Sigma-Aldrich) for 10’ at 37°, and a postfixation in 3.7% formaldehyde for 25’. The samples were then extensively washed in PBST, and then in 5X saline sodium citrate buffer containing 0.1% Tween-20 (SSCT), before being pre-hybridized in hybridization buffer (Molecular Instruments) for 1h at 37°C. The probes were then added to the hybridization buffer at a final concentration of 0.05μM and the samples were let to hybridize at 37°C overnight under gentle agitation. Following hybridization, the samples were washed 4 times 30’ in probe wash buffer (Molecular instruments) at 37°C and then in 5X SSCT at room temperature. They were then pre-amplified in amplification buffer (Molecular Instruments) for 30’. Meanwhile, H1 and H2 components of the HCR hairpins B1, B2 or B3 coupled either to Alexa546 or Alexa647 fluorophores (Molecular Instruments) were incubated separately at 95°C for 90”, cooled down to room temperature in the dark and then pooled together before being added to the amplification buffer at a final concentration of 60nM. The amplification was then performed overnight. The samples were subsequently washed 4 times 30’ in 5X SSCT and incubated in PBST supplemented in NaCl to 500mM (NaCl-PBST) and containing 1:1000 DAPI (Invitrogen) for three hours. Finally, the samples were cleared in a series of 20%, 40%, 60%, and 80% fructose diluted in NaCl-PBS. Each fructose bath was carried out for at least 1 hour. Clarified samples were mounted in 80% fructose diluted in NaCl-PBS for imaging, which was done using a Zeiss LSM700 confocal microscope. For each sample, series of optical sections were taken with a z-step interval of 2-3 μm Multichannel acquisitions were obtained by sequential imaging. Confocal optical sections spanning regions of interest along the oral-aboral axis were compiled into maximum intensity z-projections using ImageJ v.1.52g^72^.

The specificity of the antisens DNA probes and amplification hairpins was validated by running the protocol without hairpins and probes, with hairpins alone, and comparing antisens and sense probe sets for *nkx2*.*1* (Supplementary Fig. 7). The consistency of the expression patterns obtained through this method was further validated by comparisons of *elav, nkx2*.*1, dmbx* and *otx* with colorimetric whole-mount *in situ* hybridization using single antisens RNA probes as described previously^73^ (Supplementary Fig. 7).

### Immunohistochemistry

Immunofluorescence staining was performed as described previously^74^ using an anti-acetylated tubulin antibody produced in mouse (Sigma-Aldrich). For endoskeleton stainings, larvae reaching the brachiolaria stage were incubated in seawater supplemented with 5mL per 1L of saturated calcein solution (Sigma-Aldrich), a fluorescent calcium analogue that is incorporated into the endoskeleton^75^, until the completion of the metamorphosis. The post-metamorphic juveniles were then fixed, permeabilized, counter-stained with DAPI, cleared following the same procedure than for immunofluorescence stainings. Imaging was done following the same procedure than for *in situ* hybridizations.

---

**Supplementary Figure 1:**
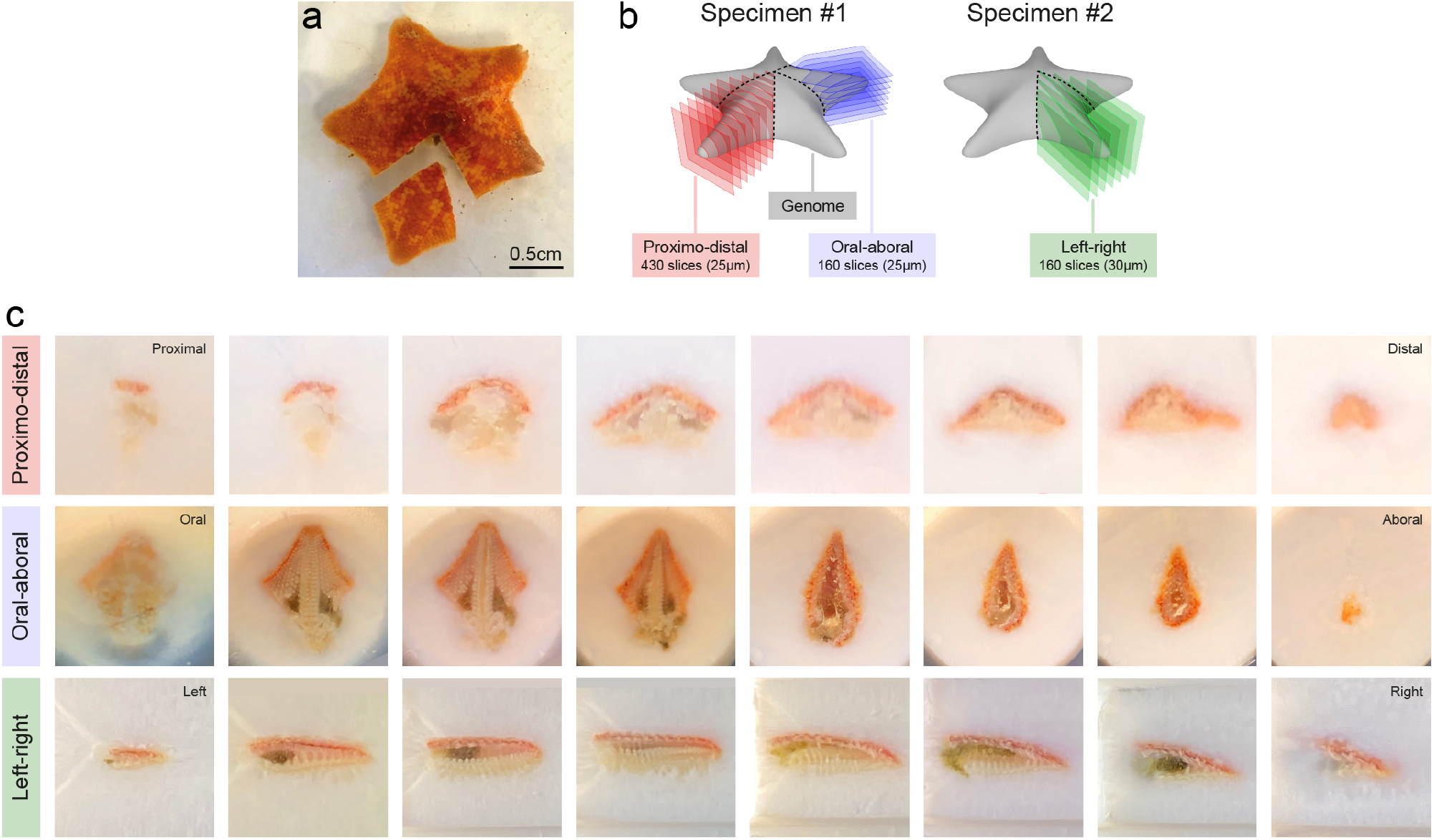
RNA tomography principle and cryosectioning. **a**, Picture of *P. miniata* specimen #1 with the first arm dissected out. **b**, Detailed experimental design of the RNA tomography. **c**, Representative pictures of through the slicing of the cryosection blocks of the P-D, O-A and L-R dimensions.

**Supplementary Figure 2:**
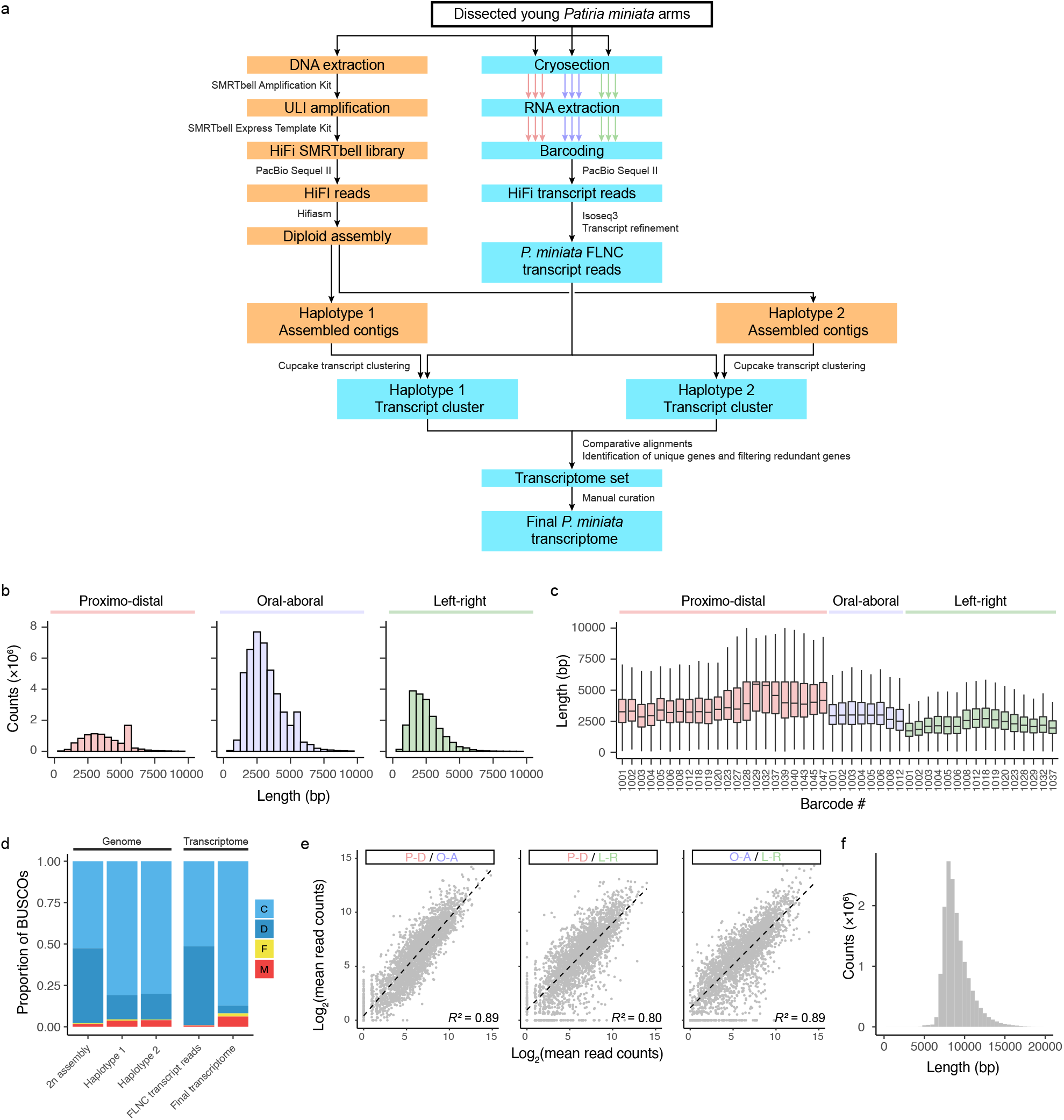
Genome sequencing and RNA tomography processing. **a**, Workflow of the RNA tomography. Orange boxes refer to genome sequencing and assembly, blue boxes refer to transcriptome sequencing and assembly. **b**, IsoSeq read length distribution in the P-D, O-A and L-R dimensions of the RNA tomography. **c**, Boxplot showing the read length per barcode in the P-D, O-A and L-R dimensions of the RNA tomography. Centre lines: median; box: interquartile range (IQR); whiskers: highest and lowest values at ± 1.5 × IQR. **d**. BUSCO assessment results for the genome (2n assembly and haplotypes) and for the transcriptome (full-length non chimeric reads and refined transcriptome). C: complete (single-copy); D: complete (duplicated); F: fragmented; M: missing. **e**, Covariance of average read counts across all sections of a dimension per transcripts show a linear correspondence of expression levels between the three dimensions of the RNA tomography. Each dot shows one of 3000 transcripts sampled out of 25,794. **f**, Genome sequencing read length density.

**Supplementary Figure 3:**
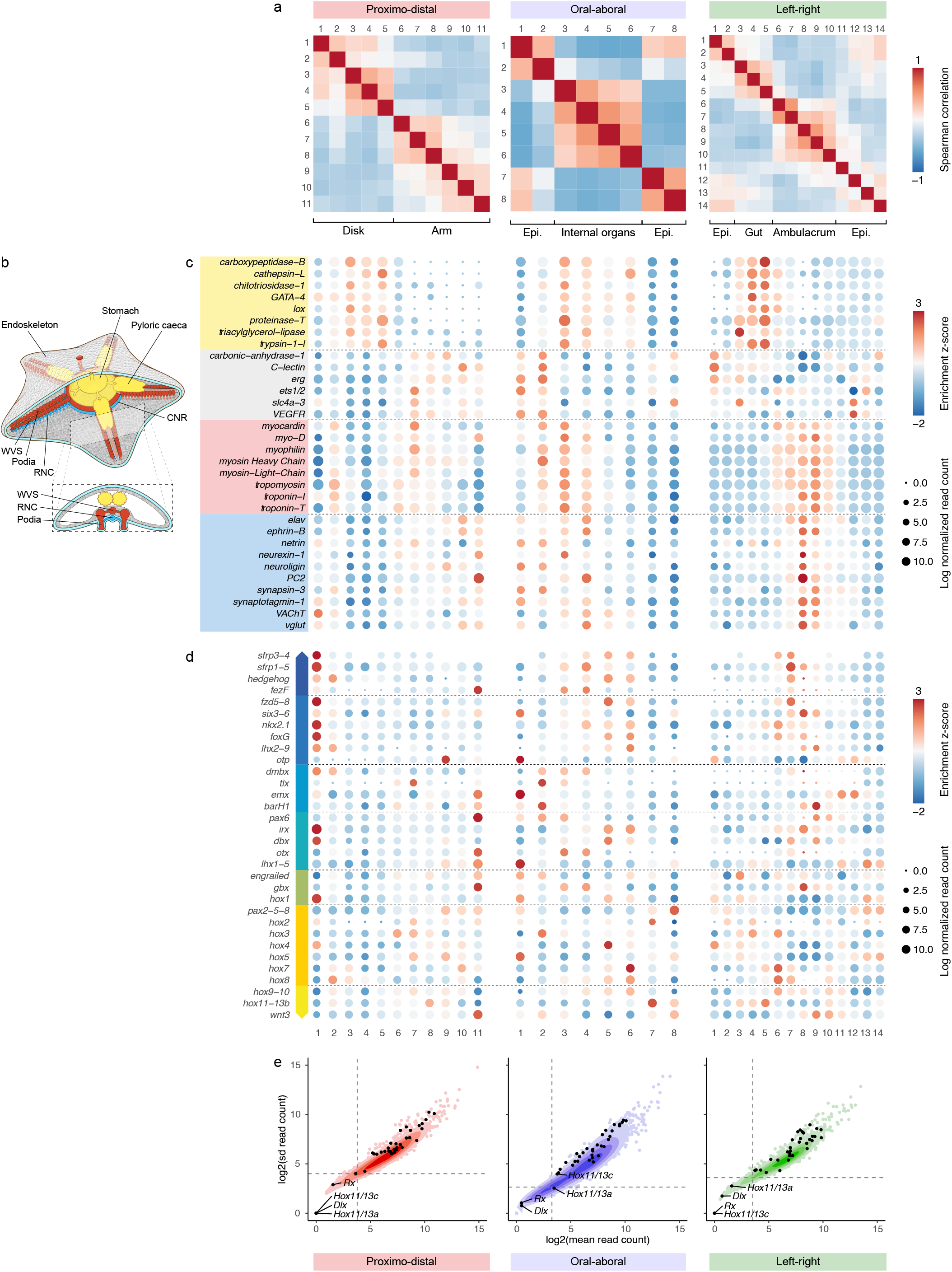
Overview of *Patiria miniata* transcriptional landscape. **a**, Spearman correlation between the sections of the three dimensions of the RNA tomography. Epi: epidermis. **b**, schematic representation of the anatomy of *P. miniata* juveniles at the stage that was used for the RNA tomography. The endoskeleton is shown in grey, endoderm, mesoderm and ectoderm derivatives in yellow, red and blue respectively. **c**, Expression profiles along the three dimensions of the RNA tomography of tissue marker genes known based on published literature to be expressed in the digestive tract (yellow), in the endoskeleton (grey), in the muscles (red) and in the nervous system (blue) are consistent with the anatomy of the animal. Note that in the case of digestive tract markers, there is a left shift of expression in the L-R dimension that we assumed to result from compression of the pyloric caeca during the dissection of the arm. **d**, Expression profiles along the three dimensions of the RNA tomography the AP patterning genes. Genes are colored according to their expression group. **e**. scatterplot showing the cutoffs (dotted lines) applied on the mean and standard deviation of the read counts in each dimension of the RNA tomography. Each dot shows one of 1000 transcripts sampled out of 25,794. AP patterning genes are shown by the black dots.

**Supplementary Figure 4:**
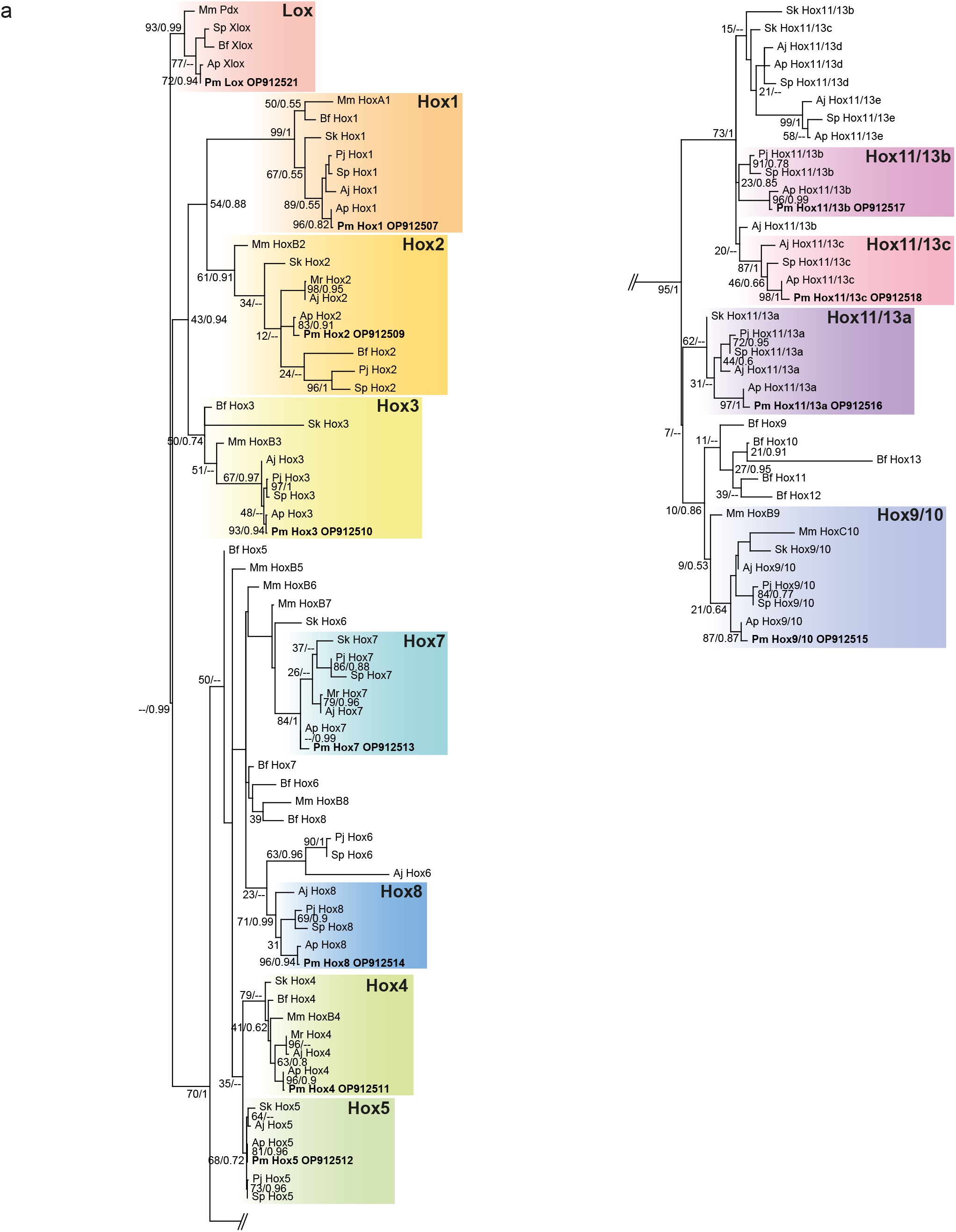
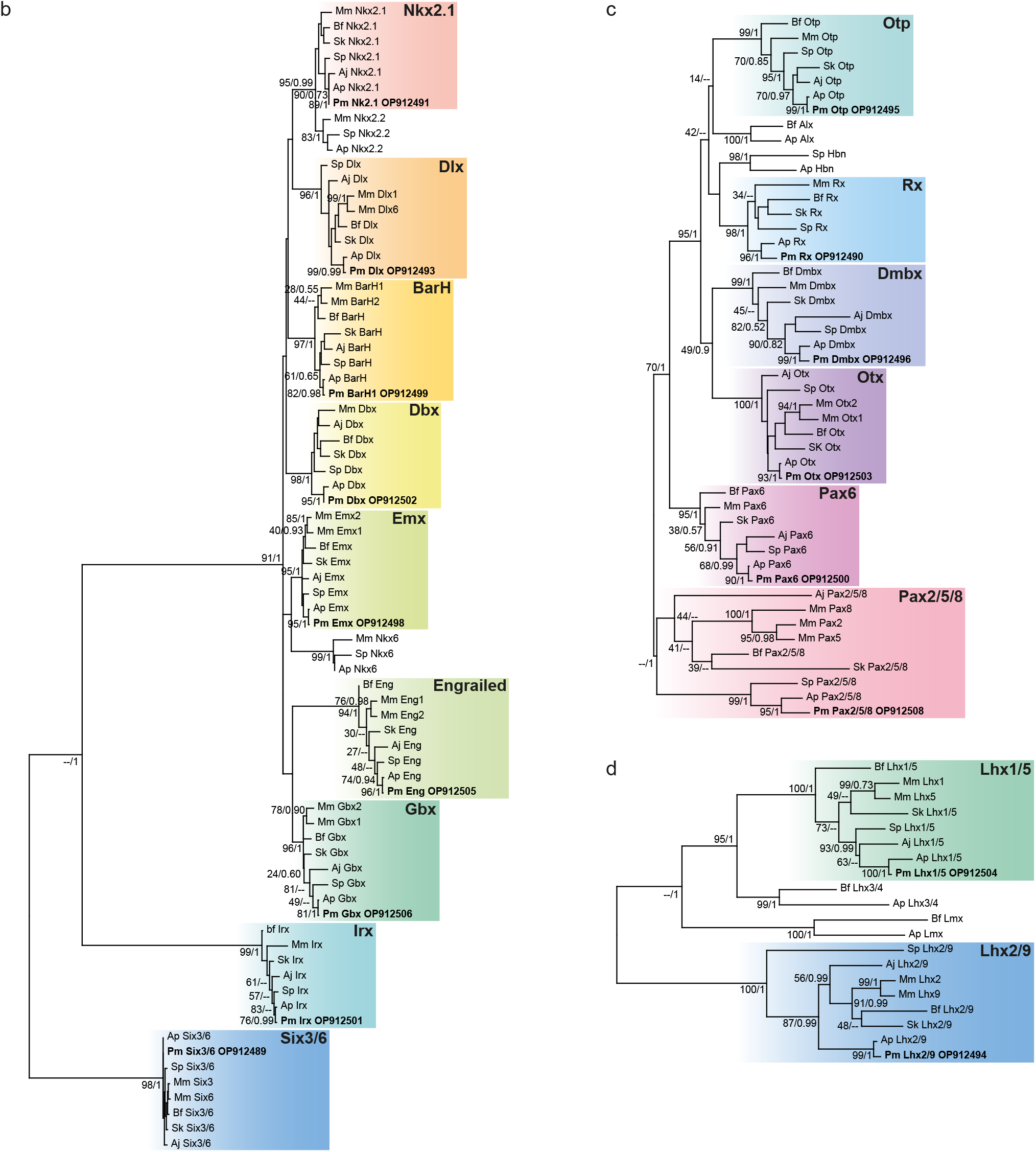
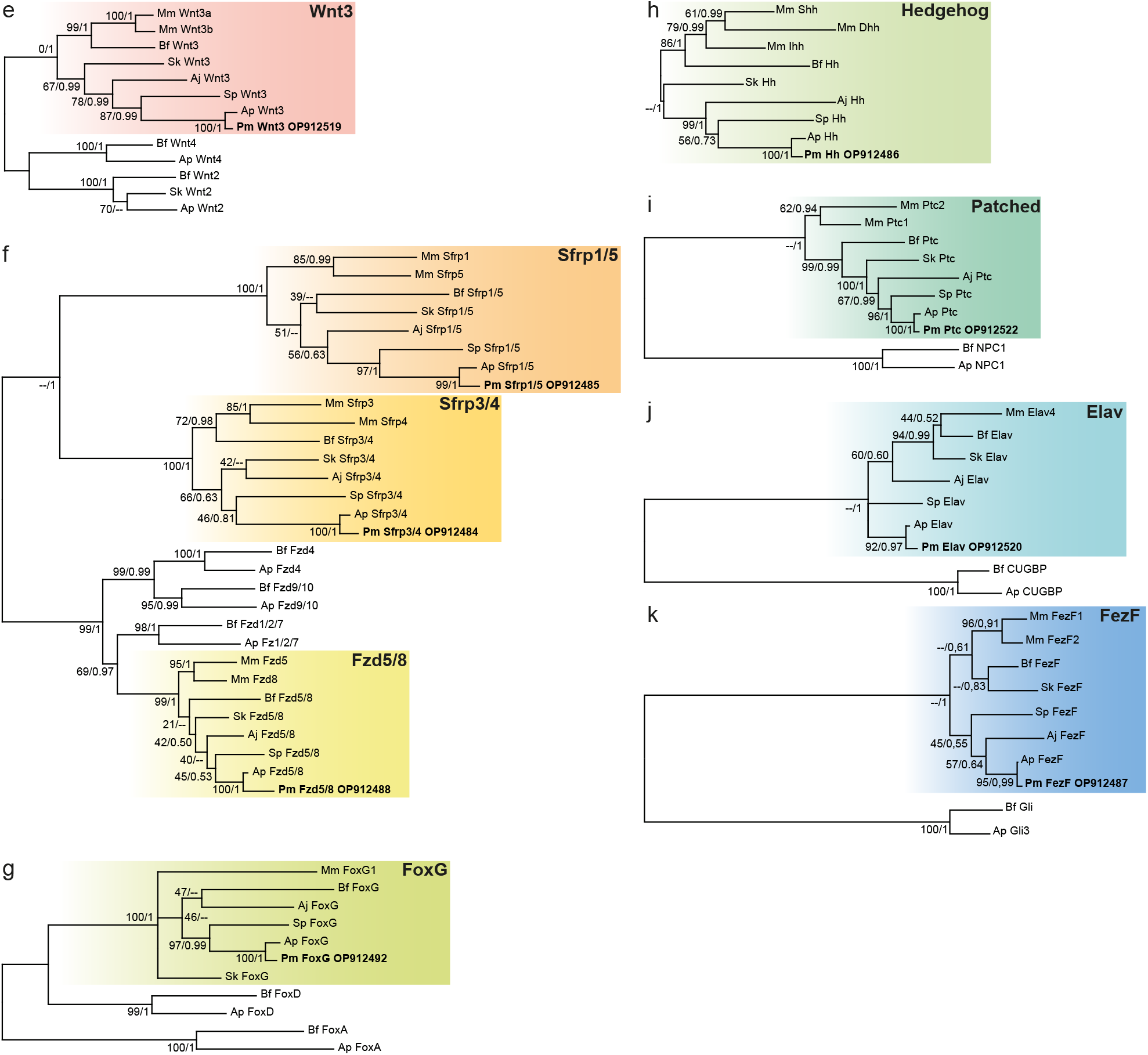
Phylogenetic trees of *Patiria miniata* orthologues. Phylogenetic relationship of *P. miniata* genes investigated in this study. **a**, Hox phylogeny. **b**, ANTP, SINE and TALE class homeobox transcription factors phylogeny. **c**, PRD class homeobox transcription factors phylogeny. d, LIM class homeobox transcription factors phylogeny. **e**, Wnt3 phylogeny. **f**, Frizzled and secreted frizzled phylogeny. **g**, FoxG phylogeny. **h**, Hedgehog phylogeny. **i**, Patched phylogeny. **j**, Elav phylogeny. **k**, FezF phylogeny. Phylogenetic trees are based on sequences from mouse (*Mus musculus*, Mm), amphioxus (*Branchiostoma floridae*, Bf), hemichordate (*Saccoglossus kowalevskii*, Sk), crinoids (*Anneissia japonica*, Aj; and *Metacrinus rotondus*, Mr), sea urchins (*Strongylocentrotus purpuratus*, Sp; and *Peronella japonica*, Pj), and crown-of-thorns sea star (*Acanthaster plancii*, Ap). *P. miniata* sequences are highlighted in bold with the DDBJ/ENA/GenBank accession number. Although trees were calculated using both the Maximum Likelihood (ML) and Bayesian Inference (BI) methods, only the ML tree is shown, with branch lengths being representative of sequence substitution rates. Branch support is indicated as bootstrap percentages from the ML analysis (ranging from 0 to 100) / posterior probabilities (ranging from 0 to 1) from the BI analysis. “--” indicates that the branching patterns of the ML and BI analyses diverged at this node.

**Supplementary Figure 5:**
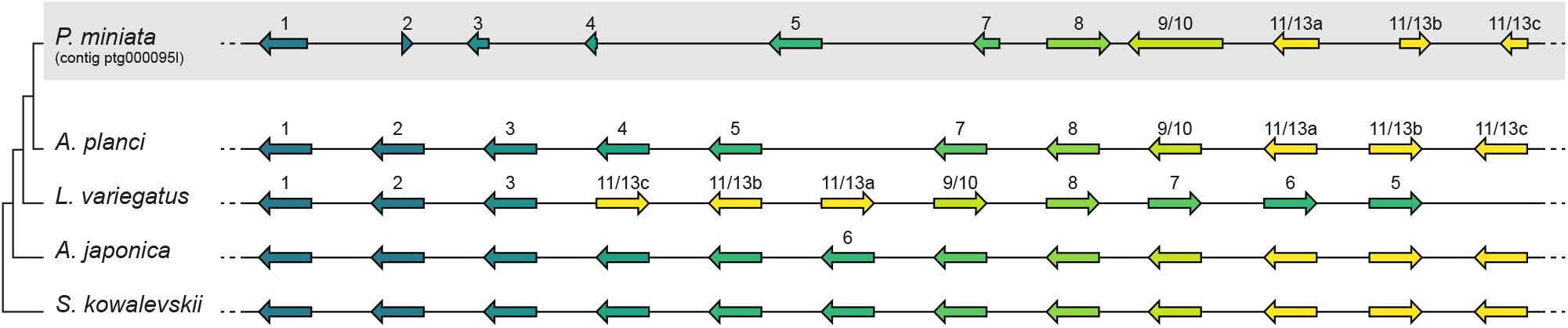
Organization of the Hox cluster in *Patiria miniata*. Representation of the Hox cluster in *P. miniata* as organized in our contig assembly. Hox clusters in *Acanthaster planci* (asteroid), *Lytechinus variegatus* (echinoid), *Anneissia japonica* (crinoid) and *Saccoglossus kowalevskii* (hemichordates) are shown for reference and are not a scale.

**Supplementary Figure 6:**
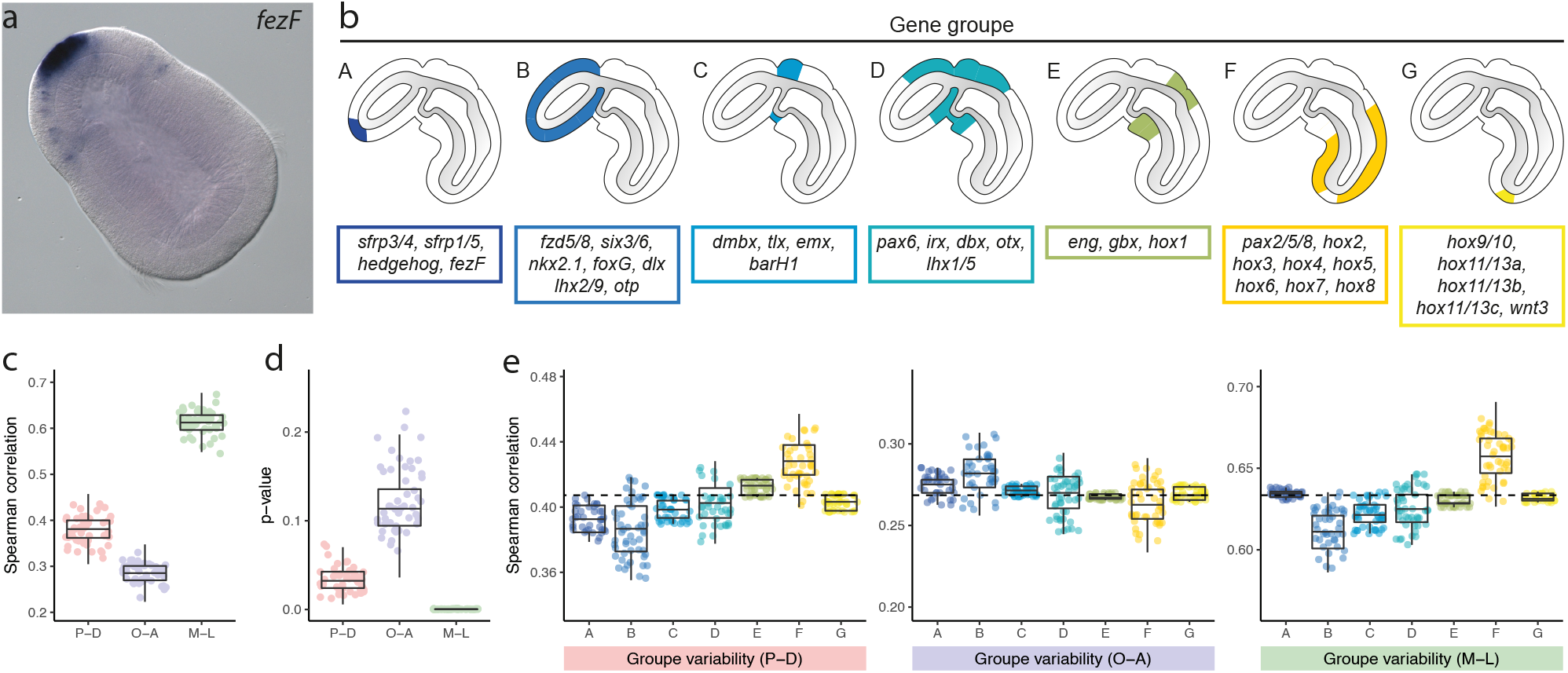
Antero-posterior gene ranking for correlation analyses. **a**, Colorimetric *in situ* hybridization for *fezF* in *Saccoglossus kowalevskii* showing expression in the anterior proboscis ectoderm and in scattered cells in the posterior proboscis ectoderm (anterior is shown at the top left of the image). **b**, Diagrams showing summarized expression patterns of AP patterning genes in *S. kowalevskii*, ranked into seven groups (A: anterior proboscis; B: proboscis; C: anterior collar; D: posterior proboscis and collar; E: collar/trunk boundary; F: trunk; G: posterior tip of the trunk) based on their anterior to posterior localization in the ectoderm. Except for *fezF*, the gene expression patterns are summarized based on data previously published (see Supplementary Table 7). **c**, Boxplot showing the distribution of the Spearman correlations between the AP gene ranking and the peak position for each dimension of the RNA tomography following 100,000 permutation of the gene ranking within each group simultaneously. **d**, Boxplot showing the distribution of the Spearman test *p*-values for the correlations between the AP gene ranking and the peak position for each dimension of the RNA tomography following 100,000 permutation of the gene ranking within each group simultaneously. **e**, Boxplots showing the distribution of the Spearman correlations between the AP gene ranking and the peak position for each dimension of the RNA tomography following 1,000 permutation of the gene ranking within each group independently. The dotted lines corresponds to the default correlation value without permutation. In **c-e**, each dot is the result of a permutation and 300 permutations sampled out of 100.000 are shown on each plot. Centre lines: median; box: interquartile range (IQR); whiskers: highest and lowest values at ± 1.5 × IQR.

**Supplementary Figure 7:**
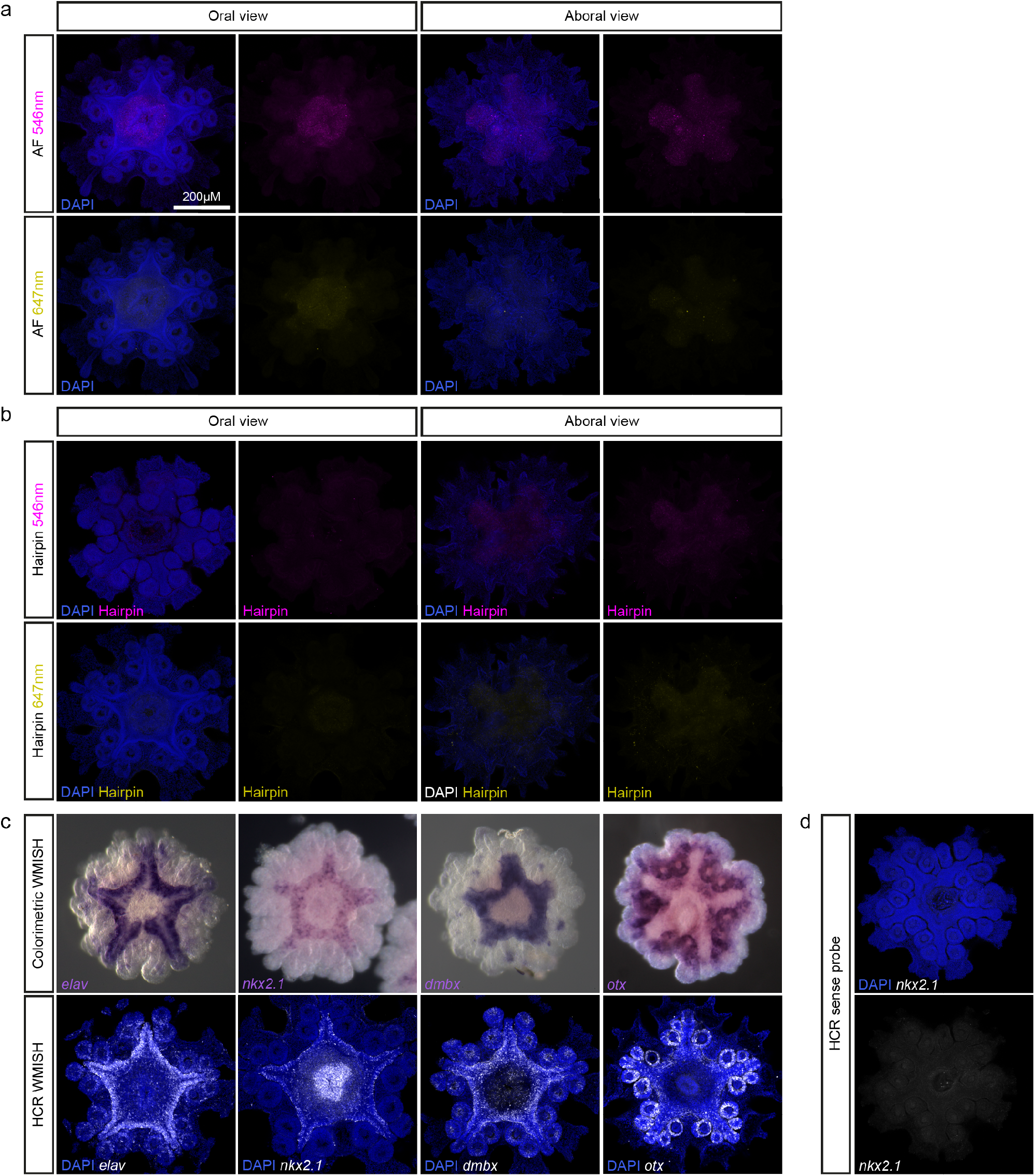
HCR control experiments. **a**, HCR controls without probes or hairpins and imaged at 546nm and 647nm, showing only the autofluorescence of *Patiria miniata* juveniles. **b**, HCR controls with Alexa546 and Alexa647 hairpins but no probes, showing the absence of non-specific binding of the hairpins in *P. miniata* juveniles. **c**, Comparison of colorimetric *in situ* hybridizations (upper row) and HCR *in situ* hybridizations (lower row) for *elav, nkx2*.*1, dmbx* and *otx* in *P. miniata* juveniles (oral side) showing that HCR demonstrates accurate and more detailed expression patterns especially in regions with low expression levels. **d**, HCR control using a sense probe for *nkx2*.*1* in *P. miniata* juveniles (oral side) showing no non-specific probe binding.

**Supplementary Figure 8:**
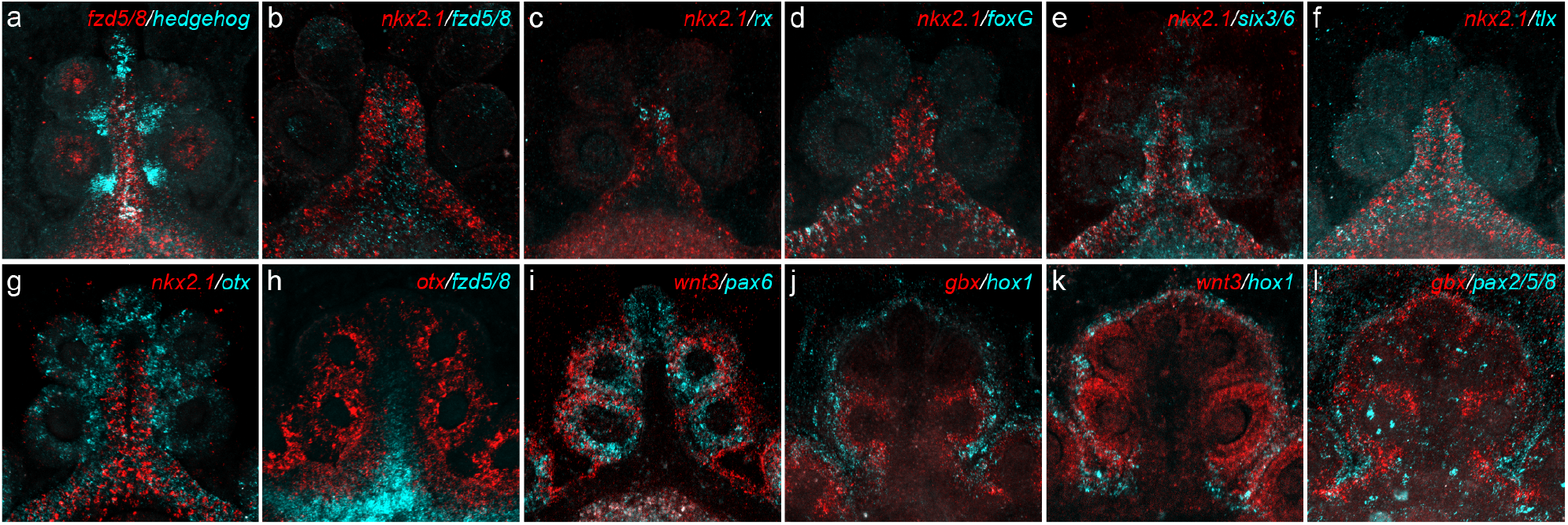
Co-expression of transcription factors in the ambulacral ectoderm. Double HCRs of *P. miniata* post-metamorphic juveniles. All pictures are showing a magnification of a single ambulacrum imaged from the oral side. **a**, *hedgehog* and *fzd5/8* colocalize along the midline of the ambulacral ectoderm, *hedgehog* is also expressed in the proximal podia epidermis. **b**, *nkx2*.*1* is expressed in all the CNR and the RNCs and outlines *fzd5/8*. **c**, *rx* and *nkx2*.*1* colocalize in the distal part of the radial nerve cords. **d**, *foxG* and *nkx2*.*1* colocalize in the circumoral nerve ring and in the proximal part of the radial nerve cords. **e**, *six3/6* and *nkx2*.*1* colocalize in the CNR and the RNCs, *six3/6* also extends in the proximal podia epidermis. **f**, *tlx* and *nkx2*.*1* colocalize in the CNR and the RNCs, *tlx* is also expressed at the tip of the primary podia epidermis. **g**, *otx* and *nkx2*.*1* colocalize on the edges of the medial ambulacral ectoderm, *otx* extends more laterally in the podia epidermis. **h**, *otx* and *fzd5/8* have mutually exclusive expression domain, with *fzd5/8* being restricted to the midline of the ambulacral ectoderm while *otx* is expressed on the edges of the medial ambulacral ectoderm and in the podia epidermis. **i**, *wnt3* and *pax6* colocalize on the outer edge of the podia epidermis, *pax6* extends to the entire podia epidermis. **j**, *hox1* outlines *gbx* expression domain at the ambulacral ectoderm boundary. **k**, *wnt3* and *Hox1* partially overlaps at the ambulacral ectoderm boundary. **l**, *pax2/5/8* outlines *gbx* expression domain at the ambulacral ectoderm boundary.

**Supplementary Figure 9:**
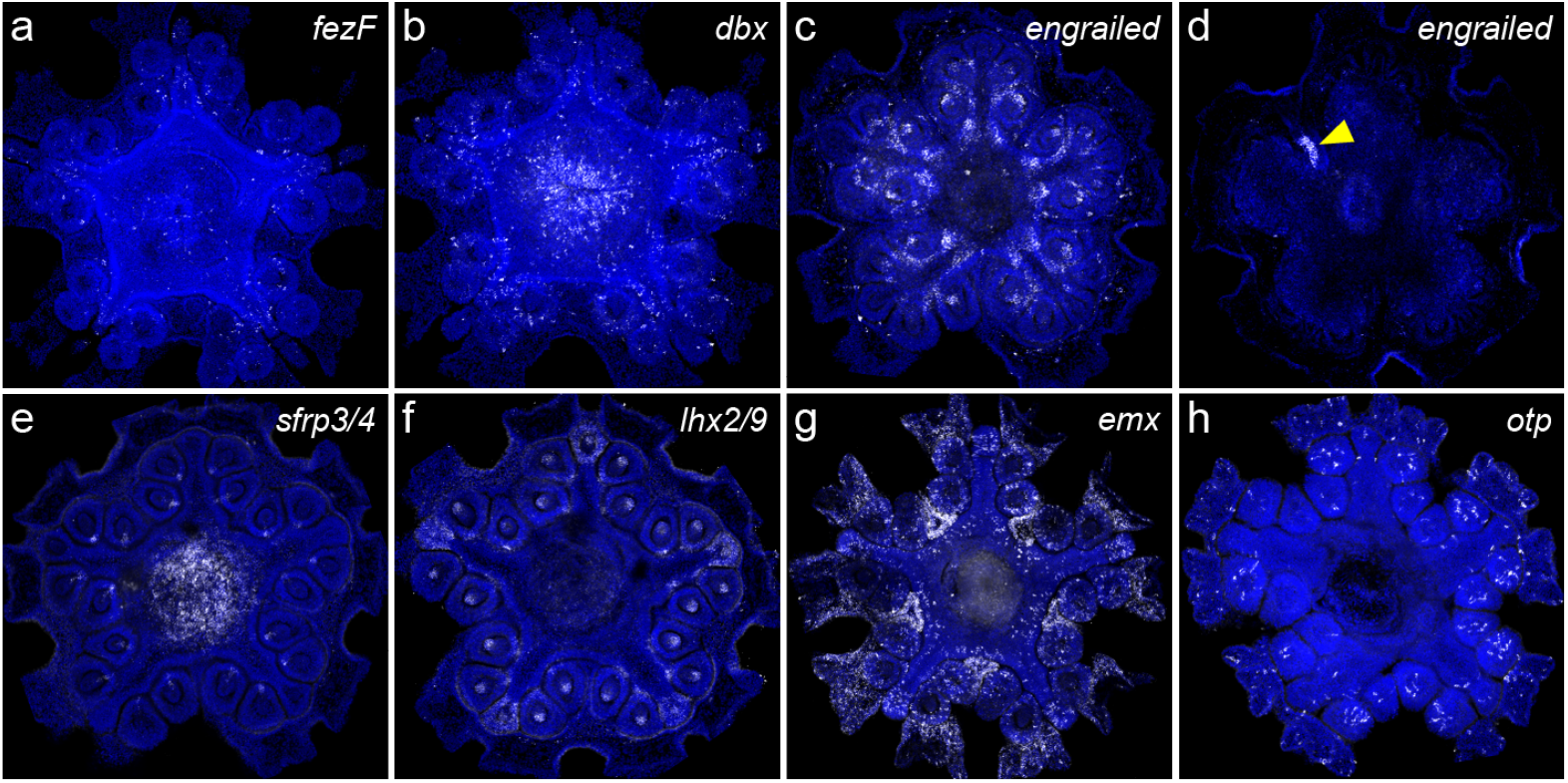
Additional expression patterns. HCRs of *P. miniata* post-metamorphic juveniles. All pictures are imaged from the oral side and are counterstained with DAPI (blue). **a**, *fezF* is expressed in scattered cells of the ambulacral ectoderms. **b**, *dbx* is expressed in scattered cells of the ambulacral ectoderm and in the pharynx. **c**, *engrailed* is expressed in the hydrocoel (radial canals). **d**, *engrailed* is also expressed in the stone canal (yellow arrow). **e**, *sfrp3/4* is expressed in the hydrocoel and in the pharynx. **f**, *lhx2/9* is expressed in the hydrocoel and in the epidermis of the primary podia. **g**, emx is expressed in scattered cells of the ambulacral ectoderm, but primarily in the ectoderm between the ambulacra and in the most peripheral regions of the interambulacral ectoderm. **h**, *otp* is expressed in scattered cells of the podia epidermis and in the interambulacral region.

